# Cas9-Assisted Biological Containment of a Genetically Engineered Human Commensal Bacterium and Genetic Elements

**DOI:** 10.1101/2021.11.03.467106

**Authors:** Naoki Hayashi, Yong Lai, Mark Mimee, Timothy K. Lu

**Affiliations:** Department of Biological Engineering, Massachusetts Institute of Technology, Cambridge, MA 02139, USA.; JSR-Keio University Medical and Chemical Innovation Center (JKiC), JSR Corp., 35 Shinanomachi, Shinjuku, Tokyo 160-8582, Japan.; Synthetic Biology Group, MIT Synthetic Biology Center, Massachusetts Institute of Technology (MIT), Cambridge, MA 02139, USA.; Research Laboratory of Electronics, MIT, Cambridge, MA 02139, USA.; Department of Microbiology, The University of Chicago, Chicago, IL 60637, USA.; Pritzker School of Molecular Engineering, The University of Chicago, Chicago, IL 60637, USA.; Broad Institute, Cambridge, MA 02139, USA.; Harvard-MIT Division of Health Sciences and Technology, Cambridge, MA 02139, USA.

**Author notes:** These authors contributed equally (Naoki Hayashi, Yong Lai).

## Abstract

Sophisticated gene circuits built by synthetic biology can enable bacteria to sense their environment and respond predictably. Biosensing bacteria can potentially probe the human gut microbiome to prevent, diagnose, or treat disease. To provide robust biocontainment for engineered bacteria, we devised a Cas9-assisted auxotrophic biocontainment system combining thymidine auxotrophy, an Engineered Riboregulator (ER) for controlled gene expression, and a CRISPR Device (CD). The CD prevents the engineered bacteria from acquiring *thyA* via horizontal gene transfer, which would disrupt the biocontainment system, and inhibits the spread of genetic elements by killing bacteria harboring the gene cassette. This system tunably controlled gene expression in the human gut commensal bacterium *Bacteroides thetaiotaomicron*, prevented escape from thymidine auxotrophy, and blocked transgene dissemination for at least 10 days. These capabilities were validated *in vitro* and *in vivo.* This biocontainment system exemplifies a powerful strategy for bringing genetically engineered microorganisms safely into biomedicine.

## Introduction

The microbiome has recently become a point of focus for biomedical research. Altered microbial communities have been associated with a multitude of diseases, including inflammatory bowel diseases^1,2^, liver disease^3^, metabolic disorders^4^, cancer^5^, and responses to COVID-19^6^ and other infections^7^. Synthetic biology offers powerful tools for manipulating the microbiome and elucidating the functions of its various components. Genetically engineered bacteria and sophisticated genetic circuits provide the means to investigate the exchange of biological signals between human and microbial cells and to eventually gain control over processes that can shield the body from diseases or overcome diseases at their onset.

Stringent regulations limit the use of genetically engineered bacteria to prevent them from being unintentionally released into the environment. The potential benefit of these organisms can be realized only once we have found ways to comply with these regulations. Thus, before the tools of synthetic biology can be applied, for example, to produce a useful protein *in situ*^8^, to stimulate immunity in response to cancer^9^, or to destroy a harmful virus in host^10^, technologies are needed to contain the genetically modified organisms and the genetic material they carry, which could find its way into other cells and alter the microbiome in unpredictable ways.

Many forms of biological containment have been devised for uncontrolled environments that do not require human monitoring or input^11, 12^. There are four main strategies for accomplishing biological containment: auxotrophy, kill switch, xenobiology, and physical containment. Auxotrophy takes advantage of engineered auxotrophic organisms, which are unable to synthesize an essential compound or to utilize an environmentally available compound required for their survival. The lack of the integral compound induces lethality outside the uncontrolled area^13^. The kill switch employs the conditional production of a toxic molecule whose gene expression is tightly controlled by an environmentally responsive element. Once engineered bacteria with the kill switch come into an undesirable environment, the switch is turned on and the bacteria are killed by the toxic molecule^14^. Xenobiology uses different nucleic acid bases, termed xenonucleic acids (XNA), as building blocks for an orthogonal chromosome^15^. The system thus uses an artificial genetic language as a genetic firewall so that the synthetic genetic elements in engineered bacteria would be nonfunctional in natural microorganisms. The physical containment traps the engineered bacteria in a space with well-designed materials, such as the multilayer hydrogel capsule. The bacteria can not escape, but still have access to nutrients and chemical inputs from outer environments^16^. These strategies have led to technologies that genetically prevent the engineered strains and the genetic elements from escaping from controlled environments.

Auxotrophy, one of the most promising approaches, has already been applied to develop genetically modified therapeutic bacteria as pharmaceutical agents^8^, such as the biocontainment of *Lactococcus lactis* by removing essential genes such as *thyA*^17^. The *thyA* gene encodes thymidylate synthase, an enzyme required for the synthesis of deoxythymidine monophosphate, which is converted to deoxythymidine triphosphate, used for DNA synthesis. Deficiency of *thyA* causes thymineless death unless thymine or thymidine is added to the growth media^17^. However, the thymidine-auxotrophic system has the potential risk that, depending on the genomic characteristics of the surrounding microbial community, the engineered organisms might acquire the essential gene from other bacteria by horizontal gene transfer (HGT)^18^ and escape their containment. Additionally, the dissemination of synthetic gene circuits to wild-type (WT) strains in nature is still a concern, even if gene flow barriers have been developed by introducing toxin-antitoxin gene systems onto artificial plasmids and bacterial chromosomes^19^. Genes on a plasmid are frequently transferred among bacteria; such transfer is currently a major cause of the prevalence of antibiotic-resistant bacteria^20^. The robustness and applicability of existing containment systems can be greatly improved to prevent genetically engineered microorganisms and synthetic genes from spreading.

*Bacteroides thetaiotaomicron* is an attractive cellular chassis for therapies in the human gut microbiota^21^. This species stably colonizes the mammalian gut and is prevalent and abundant in the human gastrointestinal tract^22, 23^. It has a broad ability to interact with a host as well as various other microbes because it ferments polysaccharides and glycans^24^ and metabolizes bile salts^25^. *Bacteroides* spp. have been reported as an effective treatment for colitis^26, 27^, and genetic tools have already been developed to manipulate these cells to sense, record, and respond to a stimulus in the mammalian intestine^28–30^. However, the many potential applications of *B. thetaiotaomicron* have been limited by the absence of a robust biocontainment system.

We have developed a novel biocontainment system in *B. thetaiotaomicron* that combines thymidine auxotrophy, an Engineered Riboregulator (ER) for controlled gene expression by the intended bacteria, and a CRISPR Device (CD) to achieve the reliable containment of engineered strains. This system can degrade an essential gene that could be transferred into the engineered bacteria from other bacteria and can also prevent engineered genetic circuits from disseminating to environmental bacteria. In addition to demonstrating their functionalities, we provide validation of the evolutionary stability of this system *in vitro* and *in vivo*.

We used a two-step methodology to construct a genetically modified, human commensal thymidine-auxotrophic bacterium bearing the functions mentioned above by introducing a gene cassette into the bacterial genome via recombination. This method, which can also be applied to a range of human gut-associated bacteria, enables the biological containment system to function stably with the controlled expression of a gene of interest (GOI), thus expanding future clinical applications of genetically engineered bacteria.

## Results

### Biological containment design

We developed a Cas9-assisted biological containment system that combines thymidine auxotrophy, an ER, and a CD to achieve controlled gene expression and biocontainment in engineered bacteria, and to prevent the release of the genetic circuit to environmental bacteria (Fig. 1). The ER controls the level of gene expression of a GOI that is naturally repressed in all bacteria other than the engineered strain. Only the genetically modified strain with an intact ER, composed of trans-activating RNA (taRNA) and cis-repressed mRNA (crRNA) containing cis-repressive sequences (CR) complementary to the taRNA, can express the GOI. Without any taRNA, the crRNA solely represses the expression of the GOI ^31^. Gene expression level is controllable by optimization of the ER structure. The CD has been reported as a sequence-specific bactericidal device that induces double-stranded breaks in a target DNA sequence^32^. This system can prevent both the acquisition of an essential gene (*thyA* gene) from environmental bacteria by degrading the gene, and the dissemination of transgenes to environmental bacteria by killing the bacteria harboring the artificial gene cassette with the CD. The containment system provides controlled expression, reducing the potential risk of loss of auxotrophy and dissemination of transgenes.

**Fig. 1.**
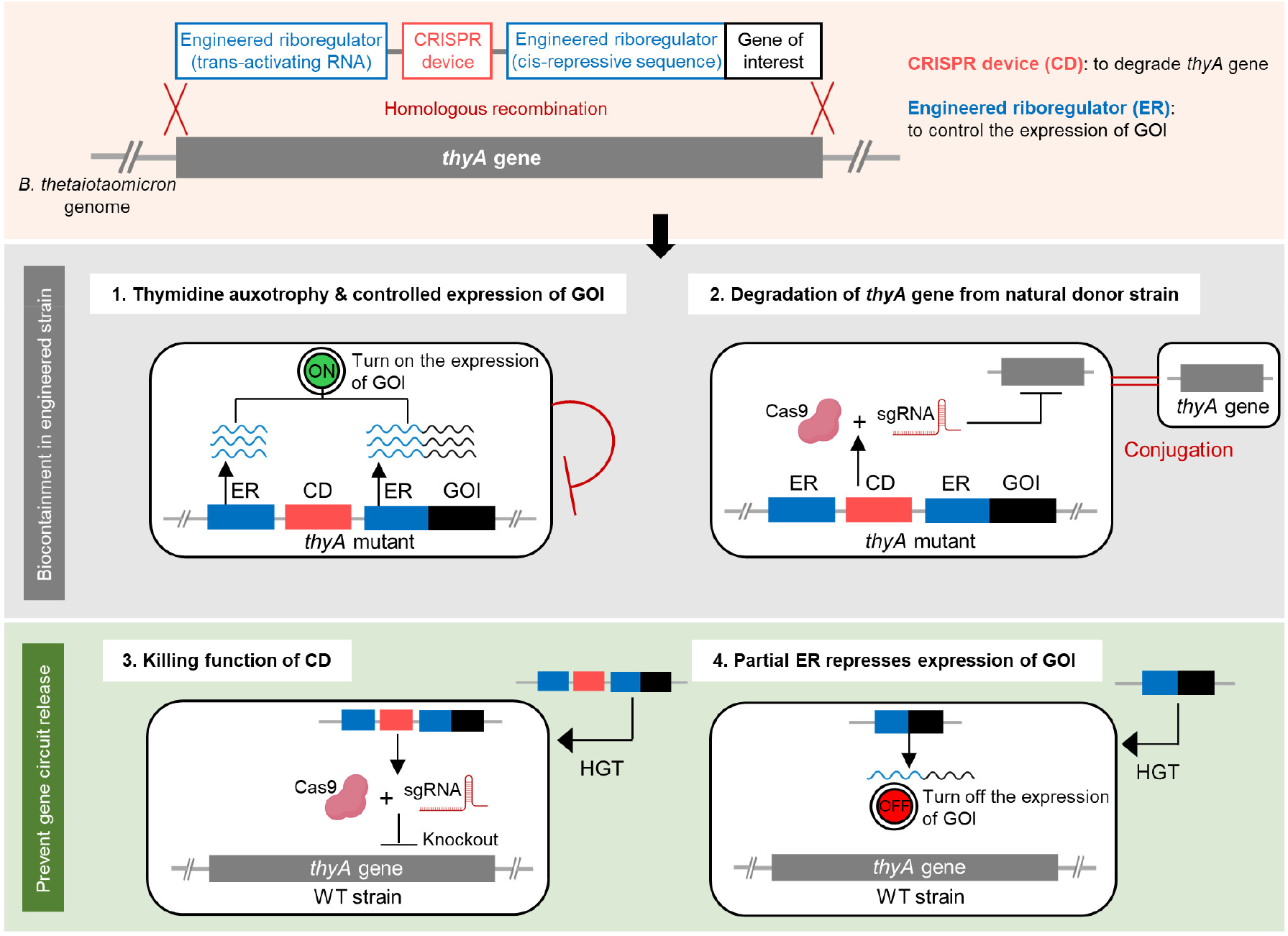
Approach for biocontainment of the genetically modified strain and genetic elements. Engineered Riboregulator (ER), CRISPR Device (CD), and Gene of Interest (GOI) are integrated into the genome of *B. thetaiotaomicron* by recombination with the *thyA* gene, which plays a role in biocontainment: **1**. Thymidine auxotrophy prevents the engineered strain from growing in the absence of thymidine. The ER controls the expression of the GOI. **2**. The CD degrades the *thyA* gene from the donor strain to maintain thymidine auxotrophy. **3**. The released CD kills the wild-type (WT) strain to prevent the dissemination of the gene circuit by knocking out the essential *thyA* gene in the genome. **4**. When released to the WT strain, partial ER represses the expression of the GOI. HGT, horizontal gene transfer.

The containment system was generated through recombination of plasmids bearing the ER and the CD (Supplementary Fig. 1) into the genome of *B. thetaiotaomicron*. Upstream and downstream sequences of the *thyA* gene were introduced flanking the gene cassette, such that double crossover recombination would lead to the integration of the ER and CD and the deletion of the *thyA* gene. The taRNA, SpCas9 gene, CR, and NanoLuc gene as a reporter of the GOI were introduced to the WT *B. thetaiotaomicron* VPI-5482 strain as the first step; sgRNA to target the *thyA* gene was integrated as the second step (Supplementary Fig. 2a). We then evaluated the characteristics of NanoLuc-producing genetically modified *B. thetaitaomicron* with the containment system.

### Repression and control of gene expression by the Engineered Riboregulator (ER)

The ER is composed of two components: the crRNA and the taRNA. The secondary structure in the 5’ untranslated region (UTR) of the crRNA which has cis-repressive sequence (CR) blocks access to the ribosome binding site (RBS) and represses the expression of the downstream gene. Sequences of taRNA containing trans-activating sequence (TA) were optimized to hybridize to the crRNA and expose the RBS, initiating the translation of the NanoLuc reporter gene (Fig. 2a).

**Fig. 2.**
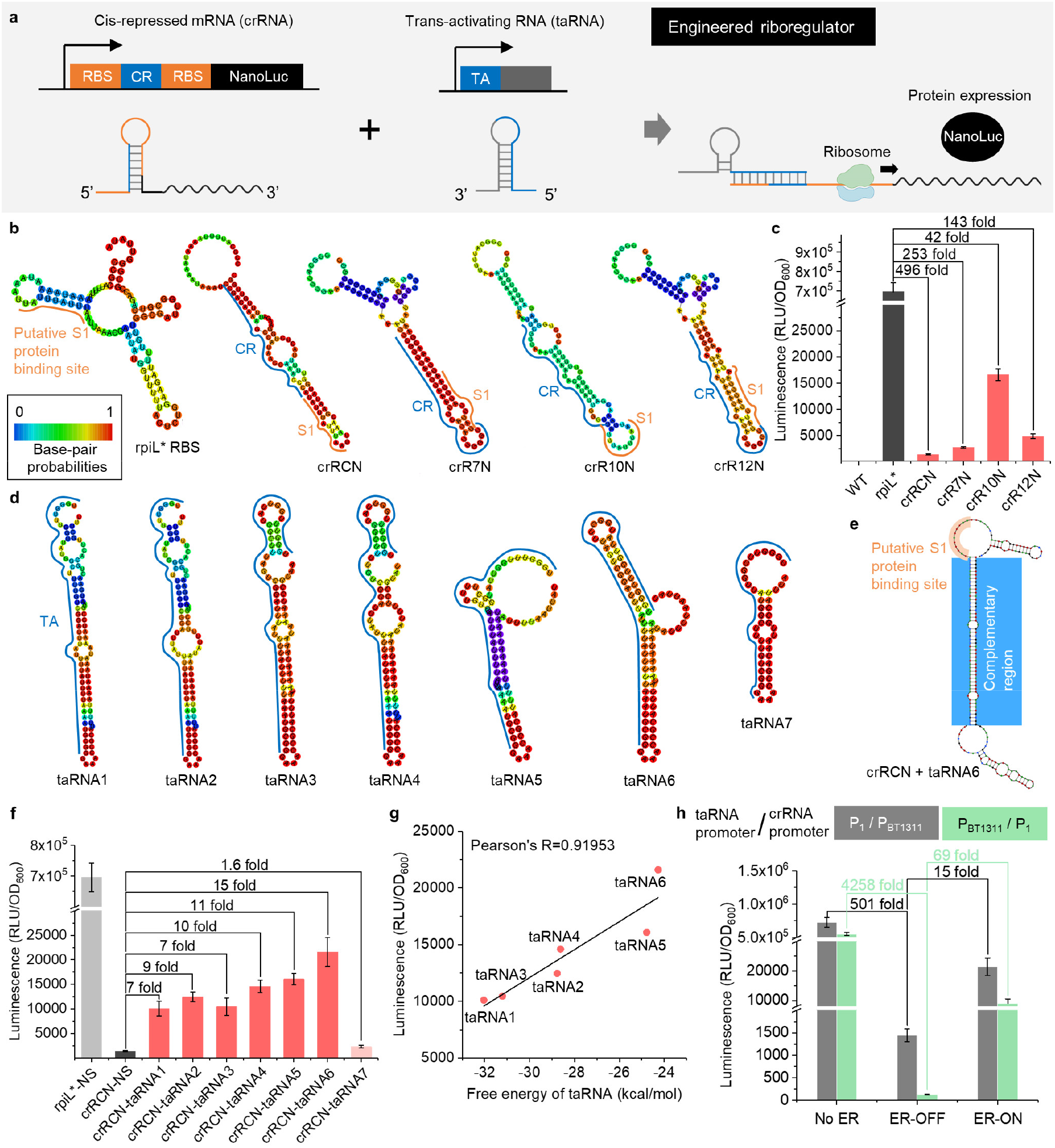
Engineered riboregulator (ER) system used to control the expression of the gene of interest in genetically modified *Bacteroides thetaiotaomicron*. **a**, Schematic of ER system. CR and TA represent cis-repressive and trans-activating sequences, respectively. **b**, RNAfold-predicted structures of cis-repressed mRNA (crRNA) variants. **c**, Results of cis repression of crRNA variants. The luminescence values were normalized to OD_600_ of cell suspensions. Fold changes were calculated on the basis of the value with no CR (rpiL*) and non-sense taRNA. **d**, RNAfold-predicted structures of trans-activating RNA (taRNA) variants. **e**, NUPACK-predicted structure of crRNA (crRCN)/ taRNA6 complex. **f**, Outputs of NanoLuc for taRNA variants. NS represents non-sense taRNA. **g**, Correlation between outputs and free energy of folding for taRNA variants: taRNA1 to taRNA6. **h**, ON/OFF outputs with P_BT1311_ and P_1_ promoters switched between crRNA and taRNA. ER-OFF, with crRCN and NS; ER-ON, with crRCN and taRNA6. Columns and dots are the values of the mean. Error bars represent the standard deviations of three biological replicates made on different days.

We first sought to build the crRNA structure to highly repress the expression of NanoLuc by inserting four types of CRs into the RBS sequence (**Supplementary Table 1**). Unlike *E. coli*, in which the homology of the Shine-Dalgarno sequence to the 16S rRNA largely dictates the strength of translation, the strength of gene expression in *Bacteroides* correlates poorly with the level of RBS complementarity to the 16S rRNA of the host organism. The RBS of *Bacteroides* tends to be AT-rich and more sensitive to secondary structure than that of *E. coli*^29^. The putative S1 protein binding site in the RBS was reported to have a strong effect on the strength of the RBS^33^. Therefore, we targeted this region to make crRNA structures. To design the ER, we used RNAfold webserver, a server designed for the analysis of nucleic acid systems^34^. RNAfold-predicted structures of crRNA were generated and showed various states of the putative S1 protein binding site with base-pair probabilities ranging from zero to one (Fig. 2b).

To evaluate the degree of repression of the GOI by crRNA, we tested the intensity of luminescence by NanoLuc. All strains except for the WT strain had luciferase activity (Fig. 2c). However, the activities of those which carried crRCN or crR7N were significantly repressed: 496-fold or 253-fold, respectively, compared to the strain without CR (rpiL*). This consequence would be caused by the secondary structures of crRNA, which block the putative S1 protein binding site and its higher base-pair probability. The rpiL* showed much higher output than crR10N even though the putative S1 protein binding site was blocked. Compared to crR10N (green color), the secondary structure of rpiL* (blue color) had the lowest probability of base-pairing, and it is possible that the rpiL* might form a different open structure in a cell.

Seven types of taRNA were then designed, by introducing inner loops and bulges to destabilize the structures, so that they had different Minimum Free Energy (MFE) and length complementary to crRNA (**Supplementary Table 2**). We thought that, in addition to the MFE, the mismatches would contribute to resistance to RNAse III cleavage of RNA duplexes^31^. The structures of taRNA were analyzed using RNAfold (Fig. 2d), and the outputs of NanoLuc were measured. NUPACK was also utilized for the crRNA/ taRNA complex. As shown in Fig. 2e and Supplementary Fig. 3a, crRNA/ taRNA complexes had exposed putative S1 protein binding sites. The crRCN and crR7N significantly repressed the expression of NanoLuc in the absence of any taRNA, while with taRNA, the NanoLuc activity of the strain that had taRNA6 (crRNA-taRNA6) was significantly increased (15 fold; Fig. 2f). The stability of taRNA affected gene expression, given that the MFE of taRNAs highly correlated with the outputs whereas the stability of the crRNA/ taRNA complex did not show any correlation with gene expression levels (Fig. 2g; Supplementary Fig. 3b). Moreover, a longer complementary sequence was required to activate the crRNA. Weaker activation of crRCN by taRNA7 was observed, a taRNA which was designed so that it would have a shorter region of complementarity (26 bps) than the others (48 bps). In contrast, the resistance to RNase III cleavage of RNA duplexes was not critical to the output, though taRNA3 was built to test resistance by adding mismatches to the RNA/ RNA complex.

To improve the ON/ OFF ratio of the gene expression level further, we switched the promoters (P_BT1311_ or P_1_) controlling taRNA and crRNA expression (Supplementary Fig. 2a). P_BT1311_ has been reported to be a stronger promotor than P ^29^. We expected gene expression to be repressed more strongly without complementary taRNA (OFF state) if the weaker promoter was placed upstream of crRNA, whereas we expected that the expression level would be retained with taRNA6 (ON state) if the stronger promoter was placed upstream of the taRNA sequence. As shown in Fig. 2h, the strain bearing the switched promoters (P_BT1311_/ P_1_) had a remarkably improved ON/OFF ratio. This taRNA elicited a ∼70-fold change in gene expression between the ON and OFF states.

### Effect of CRISPR Device (CD) on cell growth

We next investigated whether the CD, composed of the SpCas9 gene and sgRNA, affected the cell growth of the genetically modified thymidine-auxotrophic *B. thetaiotaomicron* strain. The two sets of sgRNAs and promoter sequences (P_cfxA_-sgRNA1 and P_cepA_-sgRNA1) were introduced, via the second double crossover, into the strain having crRCN and taRNA6 as an ER (Supplementary Fig. 2). P_cfxA_-sgRNA1 targets the *thyA* gene with the stronger promoter P_cfxA_. P_cepA_-sgRNA1 has a weaker promoter than the P_cfxA_-sgRNA1^29^. We next tested the growth curves of the strains in the presence and absence of thymidine. With thymidine, the P_cfxA_-sgRNA1 strain grew more slowly than the other strains (Fig. 3a), possibly because the high concentration of sgRNA was harmful to this strain. In fact, the P_cfxA_ promoter is approximately 10-fold stronger than the P_cepA_ promoter. The high concentration of the sgRNA might have increased the frequency of small-RNA-mediated targeting of mRNA^35, 36^. On the other hand, we also observed that with thymidine the P_cepA_-sgRNA1 strain grew rapidly, like the WT strain, but without thymidine this strain grew just a little and then died; the initial growth would have been enabled by the thymidine retained in the cells before the assay began. Overall, high expression of sgRNA can lead to toxicity and inhibition of cell growth.

**Fig. 3.**
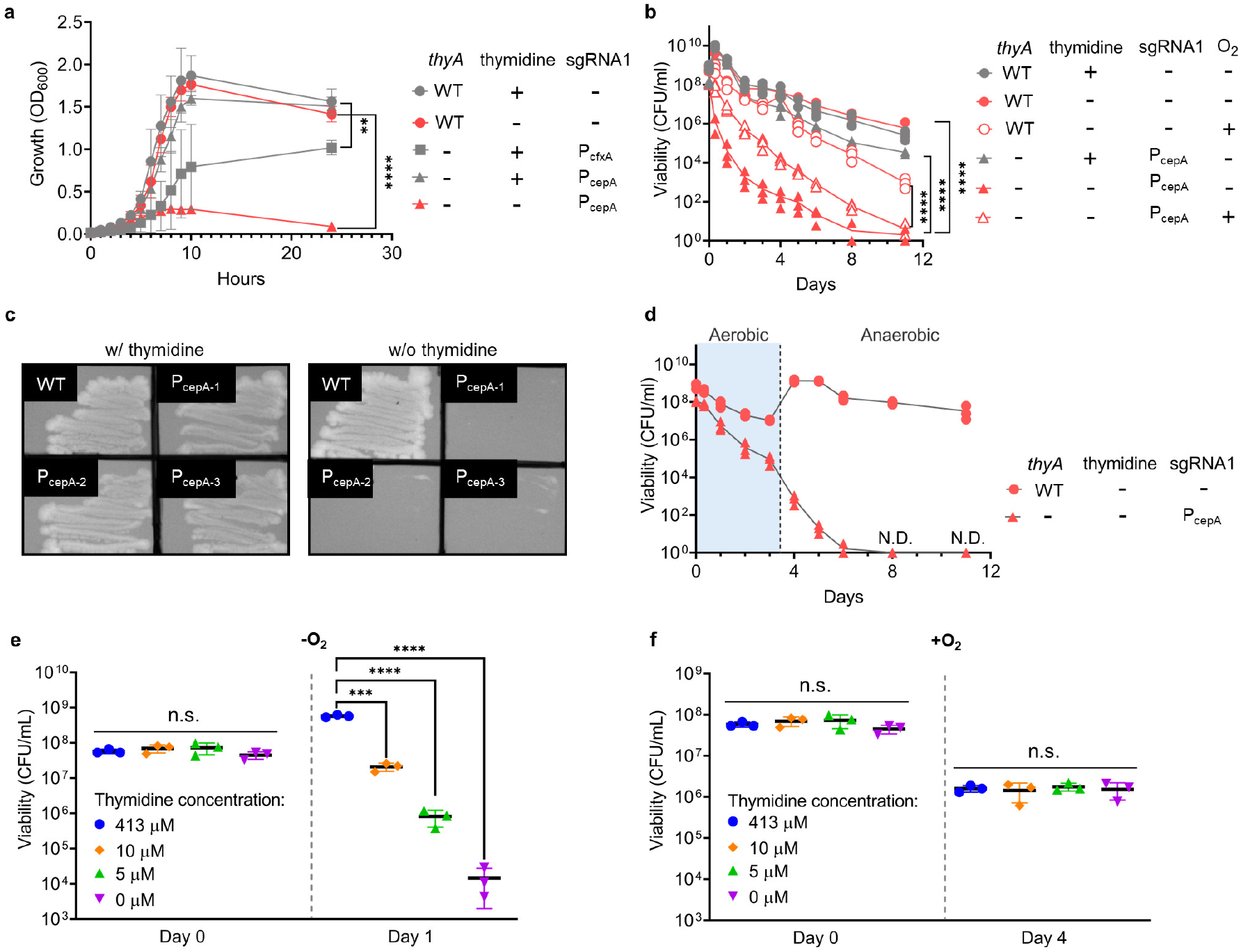
Thymidine auxotrophy of genetically modified *B. thetaiotaomicron.* **a**, Growth curves of wild-type (WT) and *thyA*-deficient strains with P_cfxA_-sgRNA1 and P_cepA_-sgRNA1 in the presence and absence of thymidine. **b**, Cell viability of WT and *thyA*-deficient strain with P_cepA_-sgRNA1 in the presence and absence of thymidine under anaerobic and aerobic conditions. **c**, Restreaked colonies of the *thyA*-deficient strain with P_cepA_-sgRNA1 on TYG with and without thymidine, after anaerobic culturing without thymidine for 11 days. **d**, Cell viability of WT and *thyA*-deficient strain with P_cepA_-sgRNA1 when cells were transferred from aerobic to anaerobic conditions in the absence of thymidine. **e**, Cell viability of *thyA*-deficient strain with P_cepA_-sgRNA1 at various concentrations of thymidine in the anaerobic condition. **f**, Cell viability of the *thyA*-deficient strain with P_cepA_-sgRNA1 at various concentration of thymidine in the aerobic condition. (**a**) Each dot is the value of the mean, and error bars represent the standard deviations of three biological replicates made on different days (Tukey’s multiple comparisons test with one-way analysis of variance on data at 24hrs, ** P<0.01 **** P<0.0001). (**b**-**f**) Each dot is a biological replicate, and lines and central bars are the values of the mean. Error bars represent standard deviations (Tukey’s multiple comparisons test with one-way analysis of variance on log-transformed data, *** P<0.001 **** P<0.0001). The detection limit is 1 CFU/ml. n.s. represents not significant. N.D. represents no detection.

### Viability of genetically modified thymidine-auxotrophic *B. thetaiotaomicron*

To validate thymineless death as a method of biological containment of the genetically modified strain, we measured viability aerobically and anaerobically. The cell concentration of the WT strain increased over 1 × 10^9^ CFU/mL within 8 hrs anaerobically and then gradually decreased as a result of lack of nutrition or the accumulation of waste products. However, the cell concentration remained higher than 1 ×10^5^ CFU/mL for 11 days. Likewise, the P_cepA_-sgRNA1 strain with thymidine showed large numbers of living cells throughout the culture (Fig. 3b). On the other hand, we observed an approximately 10^4^-fold decrease in CFU/mL of the genetically modified strain without any sgRNA and a 10^6^-fold decrease in the P_cepA_-sgRNA1 strain 6 days after incubation without thymidine (Supplementary Fig. 4). The CD seemed to help increase the death rate, which might be derived from its toxicity. When the culturing was prolonged up to 11 days, the CFU/mL of P_cepA_-sgRNA1 strain decreased by 10^7^-10^8^ (Fig. 3b). We streaked three colonies of the strain obtained at day 11 onto a TYG (trypticase-yeast extract-glucose) agar plate with and without thymidine to determine whether the viable cells were still thymidine auxotrophs. As shown in Fig. 3c, these cells showed thymidine auxotrophy and would die without any growth even if the viability test was extended after day 11.

When the P_cepA_-sgRNA1 strain was cultured aerobically, the rate of decrease in viable cells was repressed (Fig. 3b). The cell growth, or DNA replication, of this anaerobe is inhibited in the aerobic state, which would have repressed the thymineless death^37^. This result was consistent with a previous report about *Bacteroides ovatus*^18^. However, our strain decreased in cell numbers more rapidly than the WT strain and did not grow any more, even if it was transferred from the aerobic state to the anaerobic state (Fig. 3b, d).

Viability was tested at various concentrations of thymidine to clarify the concentration dependency of thymineless death (Fig. 3e, f). The decrease in viable cells was not caused by thymineless death in the aerobic state, but possibly by stress from the sgRNA and/or oxygen. On the other hand, this strain died rapidly at concentrations of thymidine less than 5 μM in the anaerobic state.

### Blocking acquisition of *thyA* gene by genetically engineered bacteria

Next, we tested whether the CD could prevent the genetically engineered strain from acquiring the *thyA* gene from another bacterial strain. When the *E. coli* S17-1 λ pir strain, carrying a plasmid bearing the intact or mutated *thyA* gene, was mixed with the genetically modified *B. thetaiotaomicron* strain, the plasmid transferred to the *Bacteroides* strain by conjugation (Fig. 4a).

**Fig. 4.**
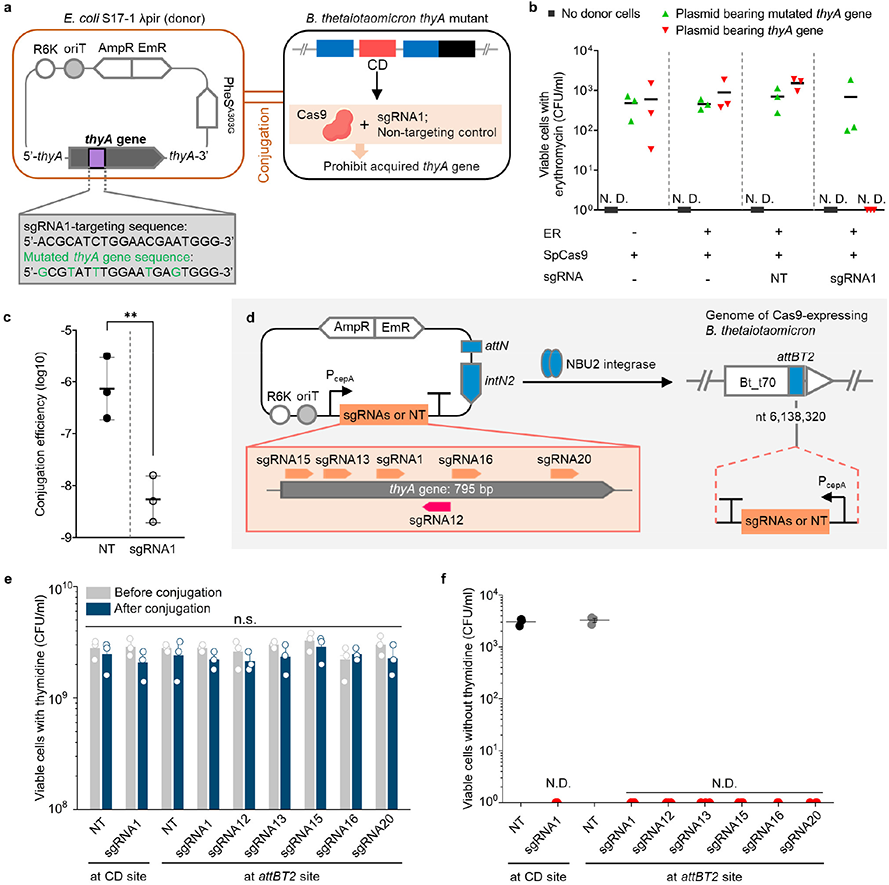
Prohibition of acquisition of *thyA* gene by genetically engineered strain. **a**, Schematic of plasmid construct bearing *thyA* gene and Horizontal Gene Transfer (HGT) of *thyA* gene through conjugation between *E. coli* and genetically engineered thymidine-auxotrophic *B. thetaiotaomicron*. The *E. coli* strains have either an intact or a mutated *thyA* gene on their plasmids. The *B. thetaiotaomicron* strains have either the sgRNA1 sequence or a non-targeting control (NT). R6K, origin of replication; oriT, origin of transfer; AmpR, ampicillin resistance cassette; EmR, erythromycin resistance cassette; PheS^A303G^, mutated α−subunit of phenylalanyl-tRNA synthetase gene; *intN2*, tyrosine integrase cassette; *attN*, recombination site with *attBT2* sites on the genome of *B. thetaiotaomicron* **b**, Viable *B. thetaiotaomicron* cells following conjugation on TYG agar plates without thymidine and plating on BHIS agar plates with erythromycin. **c**, Conjugation efficiency of *thyA* gene through conjugation on TYG agar plates in the presence of thymidine. The efficiency was calculated by dividing CFU/ml on the BHIS agar plates with gentamicin, thymidine and erythromycin by total CFU/ml (two-tailed unpaired Student’s t-test on log-transformed data, ** P<0.01). **d**, Schematic of integration of a plasmid bearing each of seven types of sgRNA sequences, including NT on the genome of SpCas9-expressing *B. thetaiotaomicron* by pNBU2 technology. **e**, Cell viability of *Bacteroides* before and after conjugation between the *E. coli* and genetically engineered thymidine-auxotrophic *B. thetaiotaomicron* with various types of sgRNAs (two-tailed unpaired Student’s *t*-test on log-transformed data between before and after conjugation). **f**, Viable *B. thetaiotaomicron* when cells were plated on TYG agar plates without thymidine after conjugation on TYG agar plates with thymidine. Three biological replicates were made on the different days. Black bars and columns are the values of the mean. Error bars are the standard deviations. The detection limit is 1 CFU/ml. n.s. represents not significant. N.D. represents no detection.

To determine whether the CD carried by genetically modified strains of *B. thetaiotaomicron* would degrade the acquired *thyA* gene, several genetically modified *Bacteroides* strains with different ERs and CDs were constructed and tested for their ability to prevent the acquisition of *thyA* gene. One strain had neither ER nor sgRNA; others had the ER with or without sgRNAs, including the non-targeting control. Plasmids bearing either an intact or a mutated *thyA* gene were generated by introducing synonymous codons into the sgRNA1-targeting sequence. Filter mating between *B. thetaiotaomicron* and *E. coli* with the plasmids was performed on TYG agar plates in the absence of thymidine.

As shown in Fig. 4b, *B. thetaiotaomicron* transconjugants were detected on BHIS agar plates with erythromycin and thymidine, indicating that these strains had acquired the plasmid-borne mutated *thyA* gene that was not targeted by a CD. Additionally, the strain with the CD having the non-targeting control, which did not degrade the *thyA* gene, obtained the intact *thyA* gene and grew on the BHIS plates. In contrast, no colonies were observed of the bacterial strain having the CD that targeted the intact, plasmid-borne *thyA* gene. This result demonstrated that the CD can specifically recognize and destroy *thyA*, blocking the effect of HGT of the plasmids. The ability to disable the gene that would otherwise correct the auxotrophy after HGT has occurred is a requirement for safe biological containment.

We also conducted a conjugation experiment on TYG agar plates in the presence of thymidine. Some viable erythromycin-resistant cells having sgRNA1 were detected, in contrast to results obtained in the filter mating without thymidine. These cells would be derived from the higher numbers of viable cells during conjugation, which leads to higher numbers of transconjugants. Nevertheless, the conjugation efficiency of the plasmid having an intact *thyA* gene was significantly decreased, at least 156 fold, by the CD (Fig. 4c). The colonies selected with an erythromycin resistance gene were still thymidine auxotrophic and did not have any *thyA* gene since no amplicon was detected by PCR using primers binding to the *thyA* gene (Supplementary Fig. 5a, b). Whole genome sequencing revealed that the backbone of the plasmid bearing the intact *thyA* gene (pNH9223) had integrated downstream of the NanoLuc gene without any integration of *thyA* gene in the transconjugant (Supplementary Fig. 5c**; BioSample accession number: SAMN20797097**). Therefore, the actual frequency of *thyA* gene transfer would be less than the frequency of erythromycin resistance gene transfer. These results also indicated that the CD could also repress the acquisition of the genes adjacent to the targeted genes.

Next, we sought to clarify whether the capability to prevent the HGT of the *thyA* gene depends on the targeted sequences in the gene. Genetically modified thymidine-auxotrophic strains bearing the CD were constructed by integration of pNBU2-based plasmids with sgRNAs and the erythromycin resistance gene after the first recombination^29^; these strains carried seven types of the sgRNAs (Fig. 4d; **Supplementary Table 3**) and their promoter P_cepA_ at one of two *attBT2* sites located in the 3’-ends of the two tRNA^Ser^ genes, BT_t70 and BT_t71, on the *B. thetaiotaomicron* chromosome. These strains showed thymidine auxotrophy and antibiotic resistance when they were plated on gentamicin-containing TYG agar plates either with thymidine and erythromycin, or without either (Supplementary Fig. 6). *B. thetaiotaomicron* strains with different *thyA*-targeting sgRNAs displayed similar viability in the presence of thymidine before and after conjugation (Fig. 4e). The filter-mating experiment using thymidine auxotrophy as a selective marker revealed that all the strains had the capacity to avoid the HGT of the *thyA* gene (Fig. 4f). The choice of targeted sequence did not seem to highly affect the capability to degrade the *thyA* gene. Furthermore, the CD worked well even though the sgRNAs were integrated at a locus far from the SpCas9 gene. Thus, there seems to be no requirement to target particular sequences of the *thyA* gene in this containment system or to integrate sgRNAs at particular sites, though further research on the effect of the targeted sequence and sgRNA site on disabling the *thyA* gene is still needed.

### Prevention of dissemination of transgenes by genetically engineered bacteria

We evaluated whether the CD could block the HGT of transgenes, such as the CD, ER, and GOI, through conjugation between *E. coli* and *B. thetaiotaomicron*. The donor strain, *E. coli* S17-1 λ pir, which carries a plasmid bearing the transgenes (shown in Fig. 5a), was mixed with WT *B. thetaiotaomicron*, the recipient. The plasmids with the CD were expected to degrade the *thyA* gene on the recipient genome so that transconjugants would not survive.

**Fig. 5.**
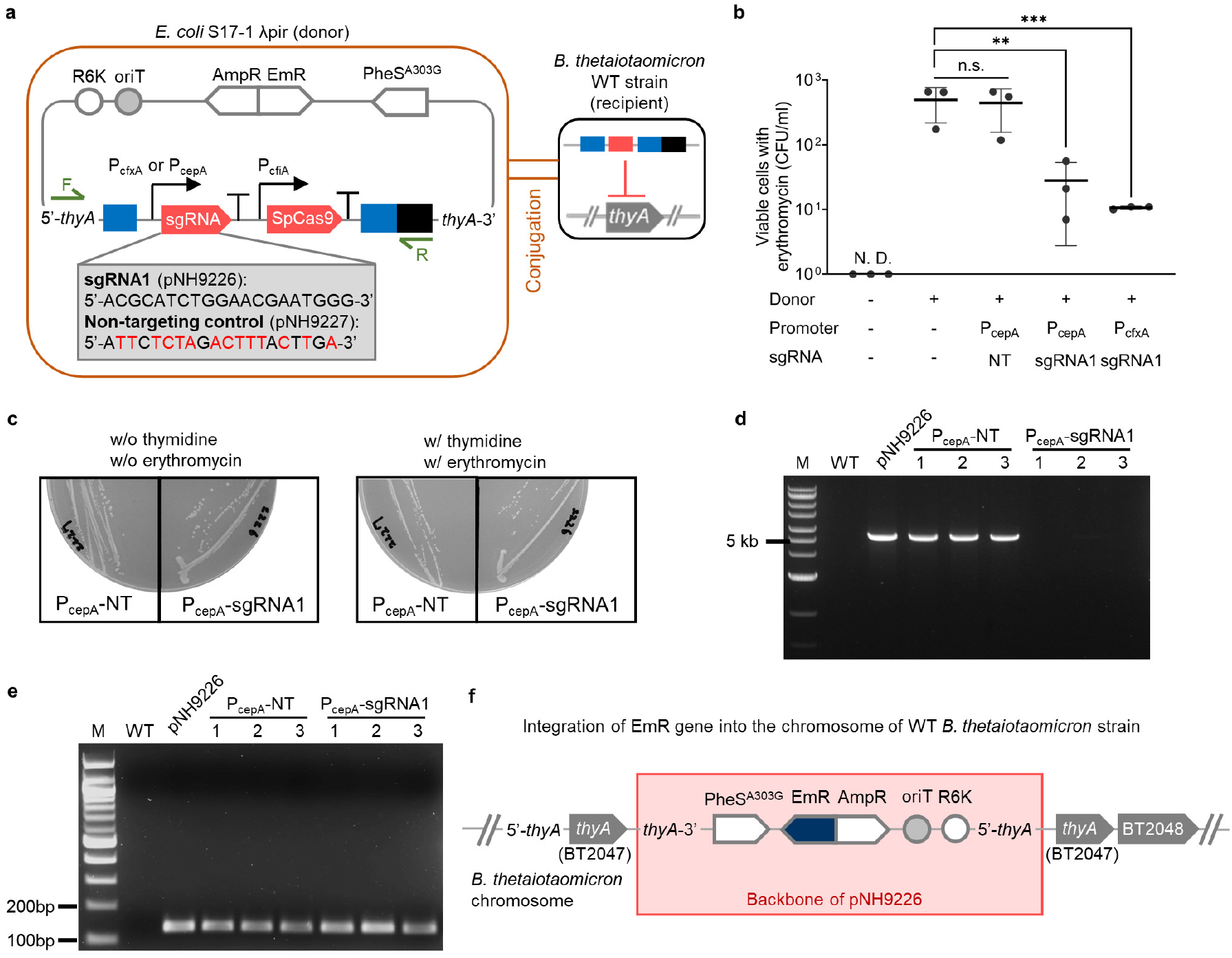
Prevention of dissemination of transgenes by genetically engineered strain. **a**, Schematic of plasmid construct bearing transgenes and Horizontal Gene Transfer (HGT) of this plasmid through conjugation between *E. coli* and wild-type (WT) *B. thetaiotaomicron*. The *E. coli* strains have either sgRNA1 or a non-targeting control (NT) on their plasmids. The *B. thetaiotaomicron* strain has an intact *thyA* gene. R6K, origin of replication; oriT, origin of transfer; AmpR, ampicillin resistance cassette; EmR, erythromycin resistance cassette; PheS^A303G^, mutated α−subunit of phenylalanyl-tRNA synthetase gene **b**, Viable *B. thetaiotaomicron* when cells were plated on BHIS agar plates with erythromycin after conjugation on TYG agar plates without thymidine. **c**, Restreaked colonies on TYG with and without thymidine and erythromycin. **d**, Amplicon patterns of three transconjugants isolated from TYG agar plates with thymidine and erythromycin after conjugation. PCR was performed with primers shown in **a**, which are named Seq-thyA-F and mmD663 in Supplementary Table 6. **e**, PCR amplification of EmR gene fragment (144 bp) in transconjugants with primers named qPCR-EmR-F and qPCR-EmR-R in Supplementary Table 6. **f**, Integration of pNH9226 backbone into the chromosome of WT *B. thetaiotaomicron* strain. Three biological replicates were made on the different days. Black bars are the values of the mean. Error bars are the standard deviations (Tukey’s multiple comparisons test with one-way analysis of variance on log-transformed data, **p < 0.01, ***p < 0.001). The detection limit is 1 CFU/ml. n.s. represents not significant. N.D. represents no detection.

When we selected the transconjugants on BHIS agar plates with gentamicin, thymidine, and erythromycin, the number of transconjugants that had received the plasmid with the CD containing P_cepA_ and P_cfxA_ was markedly lowered, by approximately 18 and 46 times, respectively, compared to that without either promoter or sgRNA (Fig. 5b). This result indicated that the *thyA* gene on the recipient genome had been destroyed by the CD and most of the recipients that had gotten the plasmid died because their genome was not repaired. The stronger the promoters utilized for the transcription of sgRNA1, the more strictly the HGT is likely to be regulated, as fewer viable cells with P_cfxA_ were detected on the BHIS agar with erythromycin than cells with P_cepA_, though the strength of the promoters did affect cell growth, as described. The appropriate choice of promoter can therefore be helpful to avoid the dissemination of the transgenes.

When the colonies of transconjugants were selected with erythromycin and restreaked on TYG in the presence or absence of both thymidine and erythromycin, they showed erythromycin resistance and thymidylate synthase activity regardless of the sgRNAs (Fig. 5c). Thus, the transconjugants with P_cep_-sgRNA1 had acquired the erythromycin resistance gene while retaining the *thyA* gene. To clarify why these transconjugants had an intact *thyA* gene after transfer of the plasmid bearing the erythromycin resistance gene and the CD, the region of transgenes in the transconjugant genomes was amplified by PCR using the primers in Fig. 5a. Interestingly, the DNA purified from transconjugants obtained by conjugation with the *E. coli* strain having P_cepA_ and sgRNA1 did not have the transgenes, whereas transgenes containing P_cepA_ and the non-targeting control were detected into the genome of WT *B. thetaiotaomicron* (Fig. 5d). As PCR amplification showed, the recipients indeed had acquired the erythromycin resistance gene (Fig. 5e). Whole genome sequencing revealed that only the backbone of the plasmid with the erythromycin resistance gene, but not the CD, was integrated into the genome of the WT recipient strain (Fig. 5f**; BioSample accession number: SAMN20797095**). The recipient may have received the erythromycin resistance gene and released the transgenes containing the CD because this portion of the transferred plasmid was detrimental to the WT strain. Therefore, the actual frequency of transgenes transferred to the recipient strains would be lower than the value discussed above.

### Evolutionary stability of genetic elements

To evaluate stability, we tested how long the ER and CD remained stable during cell growth *in vitro*. The genetically modified strain with those functions was monocultured up to 21 days and checked for gene expression of NanoLuc by the ER and the prevention of HGT of the *thyA* gene by the CD every 7 days.

Over the time course of the evolutionary stability test, the expression of NanoLuc was maintained, resulting from the stability of the ER (Fig. 6a). The genetically modified *Bacteroides* strain could receive the plasmid bearing the mutated *thyA* gene, as colonies were detected throughout the period of this experiment. On the other hand, bacterial colonies that had obtained the plasmid having an intact *thyA* gene were not observed for the first 14 days. Viable erythromycin-resistant cells appeared at day 21 (Fig. 6b). Given that there was no erythromycin-resistant colony without any donor cells (the negative control) and there was no detection of the erythromycin resistance gene by PCR in the recipients before conjugation (Supplementary Fig. 7a), we can conclude that the viable cells were not derived from spontaneous mutation during the 21 days of culture. The integration of the plasmid bearing the erythromycin resistance and the intact *thyA* genes was observed by whole genome sequencing (Supplementary Fig. 7b**; BioSample accession number: SAMN20797096**). Therefore, the CD was stable for at least 14 days under these growth condition. We assumed that some mutations may have occurred in the sequence of the CD during cell growth and that the CD may have lost functionality after day 14. However, Sanger sequencing revealed no mutation in the region of the CD for thirteen viable colonies grown on BHIS agar with gentamicin, thymidine, and erythromycin. The CD may have lost its functionality via another mechanism, such as a mutation at the locus outside the CD region, though this is still unclear. Overall, this engineered strain was stable for 14 days, which is equivalent to at least 240 cell divisions, corresponding to a 10^72^-fold amplification of the initial inoculant.

**Fig. 6.**
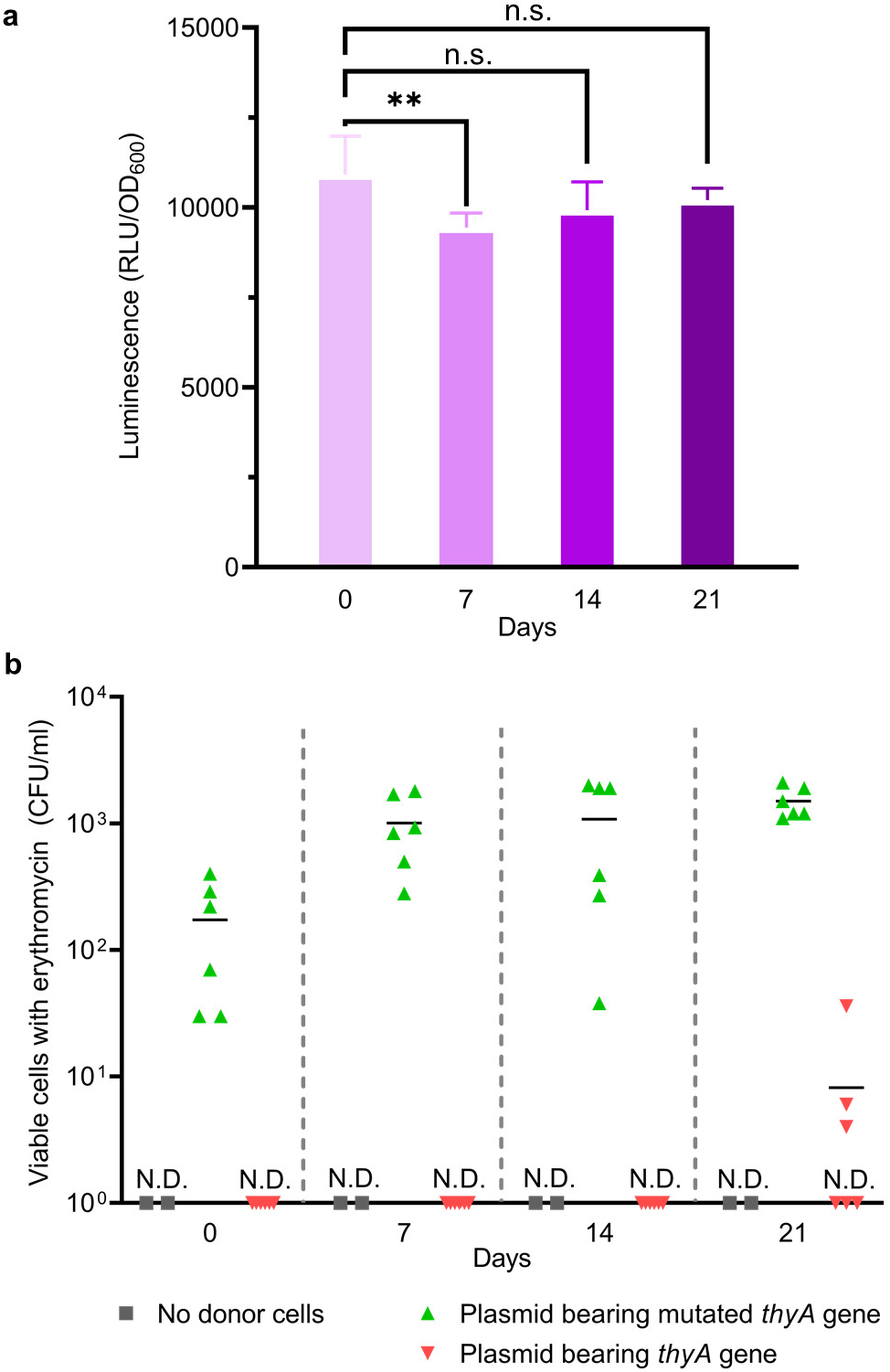
Genetic stability of the genetically engineered strain bearing Engineered Riboregulator and CRISPR Device. **a**, Outputs of NanoLuc measured every 7 days until day21. Error bars represent the standard deviations of six biological replicates (Tukey’s multiple comparisons test with one-way analysis of variance on data, ** P<0.01). **b**, Viable *B. thetaiotaomicron* when cells were plated on BHIS agar plates with thymidine, erythromycin, and gentamicin every 7 days until day21 after conjugation on TYG agar plates without thymidine. Individual dots represent six biological replicates of recipients, while two biological replicates were made for No donor cells (control). Black bars and error bars are the values of the mean and the standard deviations, respectively. The detection limit is 1 CFU/ml. n.s. represents not significant. N.D. represents no detection.

### *In vivo* colonization of the genetically modified *B. thetaiotaomicron* and function of the Engineered Riboregulator (ER) and thymidine auxotrophy

To evaluate whether the genetically modified thymidine-auxotrophic strain can colonize mice and whether the ER and thymidine auxotrophy function *in vivo*, a *thyA*-deficient strain with the ER and CD, Δ*thyA* (ER^+^/CD^+^); a *thyA*-deficient strain with the ER and the non-targeting CD, Δ*thyA* (ER^+^/CD^NT^); and a *thyA*-intact strain without either, *thyA*^+^ (ER^-^/CD^-^), were administered to mice. Stools were collected for 10 days to quantify NanoLuc luminescence and viable cells (Fig. 7a). As shown in Fig. 7b, when mice were gavaged with either the Δ*thyA* (ER^+^/CD^+^) or the Δ*thyA* (ER^+^/CD^NT^) strain after 7 days of the antibiotic treatment, the intensity of luminescence increased and then remained stable up to day 10. Thus, the genetically modified thymidine-auxotrophic strains stably colonized the mice and the ER worked well *in vivo*. High concentrations of thymidine in the mouse small intestine, which have been reported to be on the order of several hundred μM^38^, may have supported the colonization of the thymidine-auxotrophic strains. A competition experiment between the Δ*thyA* (ER^+^/CD^+^) or the *thyA*^+^ (ER^-^/CD^-^) strains was also performed. Once the *thyA*-intact strain *thyA*^+^ (ER^-^/CD^-^) colonized after the antibiotic treatment, it showed inhibition of colonization of Δ*thyA* (ER^+^/CD^+^) and the output of NanoLuc disappeared. The *in vitro* growth curve showed that *thyA*^+^ (ER^-^/CD^-^) grew faster than Δ*thyA* (ER^+^/CD^+^) (Supplementary Fig. 8). The difference in the growth rate, as well as space occupied by *thyA*^+^ (ER^-^/CD^-^) and competition for nutrients, would have inhibited stable colonization by Δ*thyA* (ER^+^/CD^+^).

**Fig. 7.**
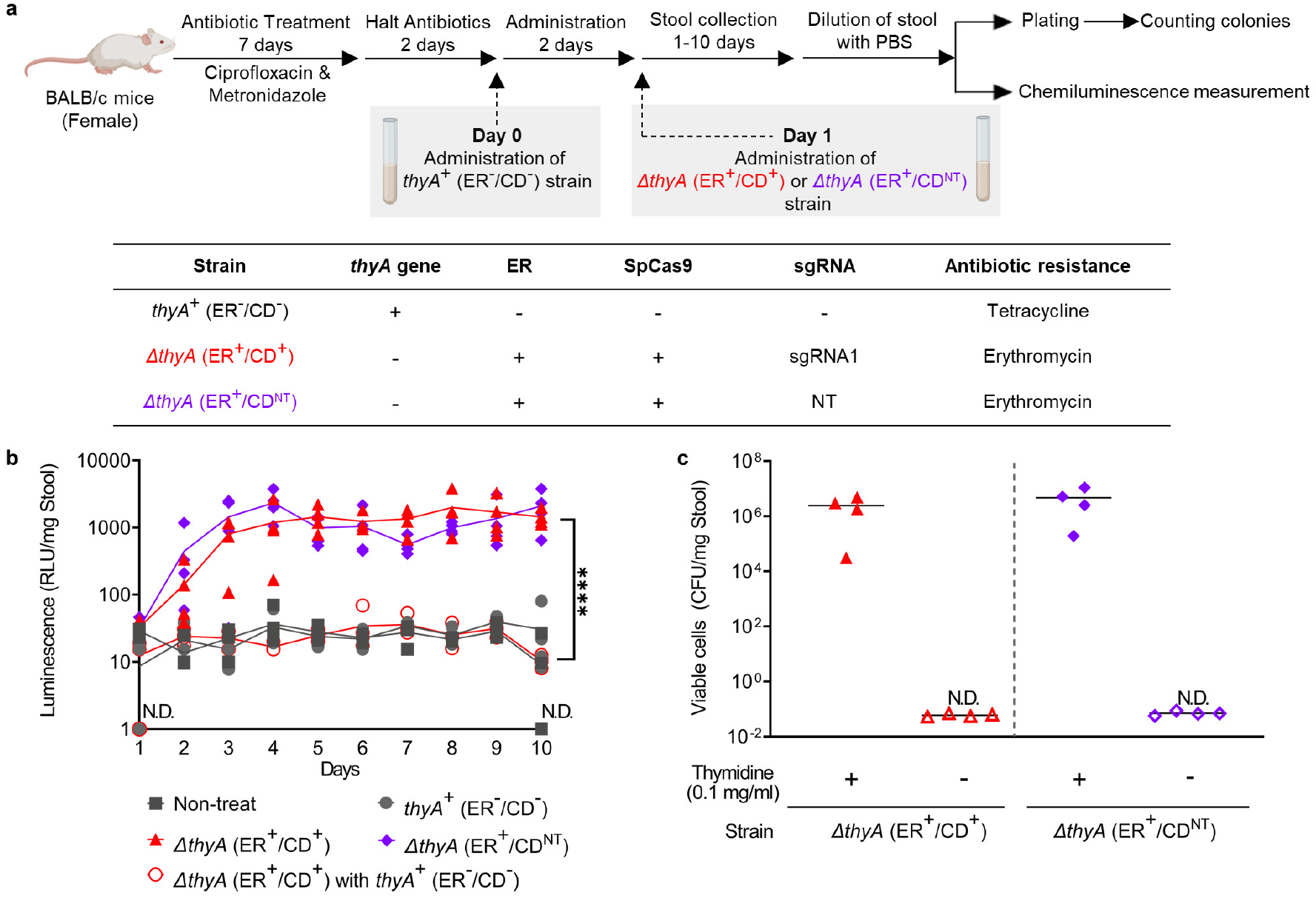
*In vivo* colonization, function of Engineered Riboregulator (ER), and thymidine auxotrophy. **a**, Experimental timeline. BALB/c mice were treated for 7 days with ciprofloxacin and metronidazole. Different *B. thetaiotaomicron* strains were administered by oral gavage on Day 0 or Day 1. The thymidine-auxotrophic *B. thetaiotaomicron* strains with ER and SpCas9 have either the sgRNA1 sequence or a non-targeting control (NT). **b**, Outputs of NanoLuc in feces of mice (gavaged with *B. thetaiotaomicron* strains as indicated) measured until day 10. Each dot represents a biological replicate, and lines are the values of the mean calculated from the four replicates (Dunnet’s T3 multiple comparisons test with Welch’s one-way analysis of variance on log-transformed data at day10, **** P<0.0001). **c**, Viable *B. thetaiotaomicron* when fecal suspensions from mice gavaged with indicated strains without *thyA*^+^ (ER^-^/CD^-^) strain were plated on selective agar plates. Black bars are the values of the mean of four biological replicates. The average detection limits are 6.0×10^-2^ CFU/mg Stool for Δ*thyA* (ER^+^/CD^+^) and 7.1×10^-2^ CFU/mg Stool for Δ*thyA* (ER^+^/CD^NT^), respectively. N.D. represents no detection.

To test the function and stability of thymidine auxotrophy *in vivo*, viable cells in fecal pellets at day 10 from the Δ*thyA* (ER^+^/CD^+^)-monoadministered and Δ*thyA* (ER^+^/CD^NT^)-monoadministered mice were enumerated by plating the fecal homogenate on selective TYG agar plates without thymidine (for cells escaping from thymidine auxotrophy) and TYG agar plates with thymidine (for total viable cells). Unexpectedly, some background growth appeared on these agar plates, but the morphology of colonies of *B. thetaiotaomicron* differed from that of the background. Therefore, we counted the colonies of the engineered strain by enumerating NanoLuc-positive colonies. Colonies escaping from thymidine auxotrophy were not detected in the fecal pellets (Fig. 7c). The average detection limits of the frequency of escape were 5.2×10^-7^ for Δ*thyA* (ER^+^/CD^+^) and 1.1×10^-7^ for Δ*thyA* (ER^+^/CD^NT^), respectively, indicating that this thymidine auxotrophy would be highly robust.

However, the number of viable thymidine-auxotrophic strains had not decreased after 7 days of the storage of solid feces at ambient temperature in the aerobic condition (Supplementary Fig. 9). This result is partially concordant with observations of *thyA*-deficient *B. ovatus* in the stools of mice^18^, in which thymineless death was repressed in the presence of oxygen. Given that the CD accelerated the death rate *in vitro*, we expected that our containment system should cause the engineered *Bacteroides* strain to decline more rapidly even in feces in the presence of oxygen than noted by Wegmann et al^18^. The genetically engineered thymidine-auxotrophic strains could have been almost completely dormant without any replication of DNA or transcription of RNA in solid feces. Another possibility is the availability of substances such as nutrients that protected the thymidine-auxotrophic strains of *Bacteroides*.

## Discussion

In this study, we describe an auxotrophic biological containment system that combines an ER and a CD. We demonstrate that this system is tunable for the expression of the GOI and highly robust, rendering the *B. thetaiotaomicron* host strain resistant to the HGT of the *thyA* gene and the dissemination of transgenes inhibited.

It has been reported that an ER could be applied to regulate levels of gene expression and create synthetic ribocomputing devices^39, 40^. An ER can also be applied as a safeguard when the taRNA and CR are designed for each individual GOI, by having the secondary structures of the crRNA include coding sequences. We created the taRNAs and CRs for the *nanoluc* gene as a model and showed the functionality of the tunable switch in a human commensal strain of *B. thetaiotaomicron*. The production of NanoLuc was switched on only when its gene was present in the cell with the CR and the corresponding taRNA. Moreover, by taking into consideration the MFE of the secondary structure when designing the taRNAs, an expression range of approximately 10-fold was obtained. A larger range of expression in the ON state and more strict suppression in the OFF state would be required for practical applications. A deep-learning approach has already been developed to optimize the structure of ERs^41^. A more sophisticated regulatory system could be created by combining the use of the ER as a safeguard and improving the method.

It has been demonstrated that auxotrophy, created by eliminating essential genes, is effective for biological containment^11, 12^. Nevertheless, it is possible that the containment system could be breached if HGT from natural microorganisms led to the acquisition of the essential genes^18^. We have constructed genetically modified thymidine-auxotrophic strains of *B. thetaiotaomicron* and quantified the loss of viable cells in the absence of thymidine. When the culturing was kept up to 11 days, the CFU/mL of PcepA-sgRNA1 strain decreased by 10^7^-10^8^. Given that the limit recommended by the National Institutes of Health^12^ for engineered microbe survival or engineered DNA transmission is less than 1 cell per 10^8^ cells, the performance of this auxotrophic system would fall just slightly short of that criterion. In particular, the death rate observed in the aerobic state was slower than that observed in the anaerobic state. The auxotrophic strain survived in the feces of mice in the presence of oxygen. Therefore, for clinical application, our system would require optimization by choosing essential genes or chassis strains to create more suitable engineered auxotrophic microorganisms for employing the ER and CD in an uncontrollable environment. Nevertheless, the CD with a promoter of appropriate strength could help to increase the death rate in the absence of thymidine whereas the engineered microorganism could grow successfully in the presence of thymidine.

The function of the CD was also verified by assessing the decrease in conjugation efficiency through filter mating. The frequency of acquisition of *thyA* gene and transgenes were suppressed by 1 to 2 orders of magnitude by the CD, which seems to be adequate to prevent the acquisition of those genes, considering the conjugation efficiency of *B. ovatus* and the target frequency described above. The conjugation efficiency of the species regarding *thyA* gene on its genome has been reported as 0.9–4.2 × 10^−7^ while the recommended efficiency is below 1 × 10^−8 18^. If the CD is utilized for gene modification, the efficiency would satisfy the criterion. The capability of the CD could be optimized by adjusting the strength of promoters, as shown in Fig. 5b, though the CD does affect cell growth, so there is a tradeoff. Potentially, the capability could also be improved by using Cas3 and the CasABCDE complex, which is analogous to Cas9 and enhances DNA degradation after the initial cleavage with 3’-to-5’ helicase and ssDNA exonuclease activities^32^. Thereby, functionality can be flexibly tuned to adapt to particular conditions of use.

Enzymes synthesized by the human microbiota generate many kinds of metabolites that affect host physiology, such as short chain fatty acids and bile acids^42, 43^. Genetically modified microorganisms bearing the related genes, as well as IL-10-producing *Lactococcus lactis* mentioned above, might offer a new therapeutic approach to treat diseases caused by an imbalance of these metabolites. For these organisms to be used effectively, for example, to avoid unintended protein overproduction by the producing bacteria and the spread of the introduced gene to other bacteria, the expression level of the desired proteins would have to be regulated. Therefore, the framework of this Cas9-assisted biological containment has a great potential for the practical use of genetically modified microorganisms.

We have provided proof-of-concept that the Cas9-assisted thymidine auxotrophic biocontainment system is tunable and useful to prevent the dissemination of transgenes and the escape from thymineless death, as well as to control the level of gene expression, simultaneously. The ON/OFF ratio of the gene expression and the death rate in unfavorable conditions still requires improvement to meet the standards for clinical application. One way to achieve a system that can be applied in clinical situations is to use other human commensal bacteria or essential genes, and to design the ER with deep learning. Future work can address these challenges and lead to the availability of useful genetically modified microorganisms.

## Methods

### Strains and culture conditions

*B. thetaiotaomicron* VPI-5482 (ATCC 29148) (GenBank: AE015928.1) was used for all evaluation of biological containment of genetically modified bacterial strains and genetic elements. It was grown in Brain Heart Infusion (Difco) media with 5 g/L yeast extract (Bacto) (BHIS), Trypticase-Yeast extract-Glucose (TYG) media or Defined Minimal Media (DMM) supplemented with 1 mg/L resazurin sodium salt (Alfa Aesar), 10 mg/L hemin (Sigma), 0.5 g/L cysteine hydrochloride (Sigma) and 1 mg/L Vitamin K3 (Sigma)^29^. TYG media contained: 10 g/L trypticase (BBL), 5 g/L yeast extract (Bacto), 1 g/L Na_2_CO_3_ (AMERSCO), 2 g/L glucose (Sigma), 80 mM potassium phosphate buffer (pH 7.3), 20 mg/L MgSO_4_·7H_2_O (Sigma), 400 mg/L NaHCO_3_ (Sigma), 80 mg/L NaCl (MACRON), 8 mg/L CaCl_2_ (Sigma), and 0.4 mg/L FeSO_4_·7H_2_O (Sigma). DMM media contained: 1 g/L NH_4_SO_4_ (Sigma), 1 g/L Na_2_CO_3_, 80 mM potassium phosphate buffer (pH7.3), 900 mg/L NaCl (MACRON), 20 mg/L CaCl_2_ (Sigma), 20 mg/L MgCl_2_·6H_2_O (VWR), 10 mg/L MnCl_2_·4H_2_O (Sigma), 10 mg/L CoCl_2_·6H_2_O (Sigma), 5 μg/L Vitamin B12 (Sigma), 4 mg/L FeSO_4_·7H_2_O (Sigma), and 5 g/L glucose (Sigma). Cultures were grown in an anaerobic chamber with an atmosphere of 2% H_2_ balanced with N_2_ and CO_2_ at 37°C. All media was pre-reduced overnight in anaerobic atmosphere before inoculation. Antibiotics, thymidine, 4-Chloro-DL-phenylalanine and agarose were added as appropriate: erythromycin (Sigma, 25 μg/mL), gentamicin sulfate (TEKnova, 200 μg/mL), carbenicillin disodium salt (Gold Biotechnology, 5 μg/mL), tetracycline hydrochloride (IBI Scientific, 5 μg/mL), Polymyxin B sulfate (Merck Millipore, 10 μg/mL), thymidine (Alfa Aesar, 100 μg/mL), 4-Chloro-DL-phenylalanine (Sigma, 2 mg/mL), and agarose (LabExpress, 15 g/L)

*E. coli* S17-1 λ pir^44^ was utilized for the construction of plasmids and the transformation of *B. thetaiotaomicron* strains through conjugation. *E. coli* S17-1 λ pir was routinely grown in Luria Bertani (LB) broth (LabExpress) supplemented with 100 μg/mL carbenicillin (Gold Biotechnology) for plasmid selection.

### Plasmid construction

All plasmids were constructed with standard cloning procedures. Briefly, DNA fragments were generated by PCR or purchased from Integrated DNA Technologies, Inc. Fragments were assembled by Gibson Assembly Cloning Method into plasmids^45^. Plasmid sequences were confirmed by Sanger sequencing. *E. coli* S17-1 λ pir was transformed with the constructed plasmids by electroporation to propagate and for transformation of *B. thetaiotaomicron* through conjugation. Genetic parts, plasmids, and primers used in this study are shown in **Supplementary Tables 1-6**. The representative plasmids are depicted in Supplementary Fig. 1: pNH9188 was used for the first recombination and pNH9196 was used for the second recombination to construct the genetically engineered *B. thetaiotaomicron* evaluated by the *in vitro* and *in vivo* study.

### Strain construction

Thymidine-auxotrophic genetically modified strains were generated through recombination of plasmids bearing ER and CD (Supplementary Fig. 1) into the genome of *Bacteroides thetaiotaomicron*. The wild-type *B. thetaiotaomicron* VPI-5482 strain was transformed with the plasmids to build genetically modified *thyA*-deficient strains through double crossovers upstream and downstream of the *thyA* locus (Supplementary Fig. 2)^46^. The taRNA sequences, SpCas9 gene, CR, and NanoLuc gene were integrated on the genome of the *B. thetaiotaomicron* by the first recombination, exchanging the *thyA* gene on the genome with transgenes. sgRNA to target the *thyA* gene was integrated by the second recombination (Supplementary Fig. 2a). The plasmids bearing those sequences were used to transform WT *B. thetaiotaomicron* through conjugation with *E. coli* S17-1 λ pir.

Overnight cultures of the WT strain were added to the *E. coli* S17-1 λ pir with the plasmids and the mating mixtures were pelleted, resuspended in BHIS medium, spotted onto BHIS agar plates and incubated upright at 37℃ aerobically. After overnight incubation, cells were collected by scraping, resuspended, and plated on BHIS agar plates with gentamicin and erythromycin as selective markers and incubated anaerobically for 2 days at 37℃. Resultant colonies were re-isolated on the BHIS agar plates with gentamicin and erythromycin.

To obtain the *thyA*-locus-exchanged mutant, each colony was anaerobically cultured for 1 day in 3 mL of BHIS with thymidine without antibiotics. Bacterial cells were collected by centrifugation, washed, and resuspended in PBS, and the appropriate dilution was plated onto DMM agar plates supplemented with p-Chloro-phenylalanine. The colonies collected from the DMM plates after 3-day cultivation at 37℃ were streaked on TYG and TYG with thymidine plates to check their sensitivity to thymidine. After 48-hour cultivation, colonies on TYG with thymidine plates were restreaked on TYG with thymidine and TYG with thymidine and erythromycin plates to check their sensitivity to erythromycin. Strains sensitive to thymidine and erythromycin were taken to be *thyA*-locus-exchanged mutants.

The genetic exchange was identified by PCR using primers which bind to the regions flanking *thyA* or to the sgRNA region. DNAs were purified from both thymidine-and erythromycin-sensitive colonies using the DNeasy Blood & Tissue Kits (Qiagen). After the DNA concentration was determined, PCR was performed using primers encompassing the exchanged site (Supplementary Fig. 2a; **Supplementary Table 6**). The amplicon sizes were compared with those from the parental strain by agarose gel electrophoresis to confirm whether the expected exchange occurred. PCR products of 2885 bp, including *thyA*, were detected in the WT strain. On the other hand, about 8000 bp of the amplicon that was bearing transgenes were detected after the double crossovers were introduced, indicating that the *thyA* gene on the chromosome had been exchanged through the recombination (Supplementary Fig. 2b).

The second recombination, i.e., that between plasmids bearing sgRNAs downstream of promoter P_cepA_ or P_cfxA_ and the chromosome of genetically modified *B. thetaiotaomicron*, was performed to integrate those sequences. The recombination process was the same as the first recombination except that thymidine was added to all of the media and agar plates. PCR was performed using primers encompassing the exchanged site and the purified DNAs as template. The presence of amplicons was checked by agarose gel electrophoresis to confirm that the expected integration had occurred. After the second recombination, the amplicons were detected at the length of about 6000 bp in strains bearing sgRNAs on the genome, whereas no amplicon was found in the parental strain (Supplementary Fig. 2c).

### Design of Engineered Riboregulator (ER)

The ERs composed of crRNA and taRNA were designed with RNAfold WebServer, which is used to predict nucleic acid structures^34^. Sequences of 99 nucleotides were simulated for crRNA; the 99-nucleotide sequences were composed of the RBS, CR, and part of the NanoLuc coding sequence, which contains the region considered to contribute to the mRNA folding effect on translation^29, 47^. Likewise, structures of 99-nucleotide sequences of taRNA were predicted. The MFE of RNA folding was predicted for each variant using default parameters.

Structures of crRNA/ taRNA complex were also analyzed with NUPACK^48^, used to predict the MFE of mRNA structures including rpiL* and part of the NanoLuc coding sequence in the previous study. NUPACK can be applied to multiple complex RNA structures whereas RNAfold analyzes a single RNA structure^34^. The MFE was calculated using default parameters other than concentrations of the crRNA and taRNA, and maximum complex size. We supposed that the concentrations of the crRNA and taRNA were estimated to be 100 nM and maximum complex size was 2^49^.

### NanoLuc luciferase assay *in vitro*

Overnight cultures in TYG medium with thymidine were diluted 1:100 in fresh medium. The cultures were grown anaerobically at 37℃ to an optical density (OD_600_) of 0.4-0.8, and the cultures were harvested by centrifugation at 4500 ×g for 10 min, washed once, and resuspended with phosphate buffered saline (PBS). NanoLuc Reaction Buffer (Promega) and cell suspensions were mixed at equal volumes. The luciferase activities were measured with an integration time of 1 second at a gain setting of 100 in BioTek Synergy H1 Hybrid Reader 6 mins after mixing. The luminescent values were normalized to OD_600_ of 300 μL of cell suspensions in PBS. Fold changes were calculated on the basis of the value with no CR (rpiL*) and non-sense taRNA, or the value with crRCN and non-sense taRNA.

### Growth curve of genetically modified thymidine-auxotrophic *B. thetaiotaomicron*

Overnight cultures in TYG medium with thymidine were harvested by centrifugation at 3200 ×g for 10 min, washed with PBS twice, and resuspended in fresh TYG medium without thymidine. The cell suspensions were diluted 1:100 in the fresh medium with or without thymidine. The cultures were grown anaerobically at 37℃ for 24 hrs. Samples (300 μL) were withdrawn from the cultures every hour for 10 hrs and at 24 hrs to monitor growth (OD_600_) in a BioTek Synergy H1 Hybrid Reader.

### Viability test of genetically modified thymidine-auxotrophic *B. thetaiotaomicron in vitro*

Overnight cultures grown in TYG with thymidine were centrifuged and washed twice in PBS, and cells were resuspended in fresh TYG media with or without thymidine, respectively. Cell suspensions were diluted in TYG with or without thymidine so that the optical density of the cell suspensions at 600 nm (OD_600_) was 0.2. Cultures were incubated anaerobically at 37°C or aerobically at ambient temperature. Inocula were withdrawn, and colony forming units were measured by plating serial dilutions on BHIS with thymidine and gentamicin in an anaerobic chamber. After incubation at 37°C for 5 days, colonies were counted manually and corrected for the dilution factor to obtain colony forming units CFU/mL.

### Assay for inhibition of the Horizontal Gene Transfer (HGT) of *thyA* gene

Overnight cultures of genetically modified thymidine-auxotrophic *B. thetaiotaomicron* strains were diluted 1:100 in fresh TYG medium with thymidine, and *E. coli* S17-1 λ pir strains bearing plasmids with the *thyA* gene and an erythromycin resistance gene were diluted 1:100 in fresh in LB medium with carbenicillin. The cultures were grown anaerobically for *B. thetaiotaomicron* and aerobically for *E. coli* at 37℃ to an optical density of 0.4-0.9 at 600nm, then harvested by centrifugation at 3200 ×g for 10 min, washed in PBS twice, and resuspended in the fresh TYG medium with or without thymidine. The optical density of the cell suspension was adjusted to 16 with the medium, and *B. thetaiotaomicron* and *E. coli* were mixed at equal volumes. Then, 50 to 100μL of the mixtures were transferred onto a 0.45 μm filter disc placed on TYG agar plates, with or without thymidine, and incubated anaerobically at 37℃ overnight. To evaluate conjugation efficiency, cells were washed off the filter in BHIS medium with gentamicin and thymidine or the TYG medium with gentamicin. The cell suspension was plated on BHIS agar plates with gentamicin, thymidine, and erythromycin or on TYG agar plates with gentamicin. Total CFU/mL of viable cells after conjugation were measured by plating the cell suspension on BHIS agar plates containing gentamicin and thymidine or TYG agar plates with gentamicin and thymidine. The colonies were counted manually 4 days after culturing.

DNA was purified from transconjugants grown on BHIS agar plates with gentamicin, thymidine, and erythromycin after conjugation, using the DNeasy Blood & Tissue Kits (Qiagen). PCR was performed with primers *thyA*-F and *thyA*-R to detect the *thyA* gene. Primers named qPCR-EmR-F and qPCR-EmR-R were used for the EmR gene (**Supplementary Table 6**). Whole genome sequencing was also performed with the purified DNA to check for the integration of the plasmid bearing the intact *thyA* gene.

### Assay for prevention of the Horizontal Gene transfer (HGT) of transgenes

Overnight cultures of WT *B. thetaiotaomicron* strains were diluted 1:100 in fresh TYG medium with thymidine, and *E. coli* S17-1 λ pir strains bearing plasmids with transgenes, such as taRNA, sgRNA, SpCas9, CR and NanoLuc gene, and an erythromycin resistance gene were diluted 1:100 in fresh LB medium with carbenicillin. The cultures were grown anaerobically for *B. thetaiotaomicron* and aerobically for *E. coli* at 37℃ to an optical density of 0.4-0.9 at 600 nm. Cultures were harvested by centrifugation at 3200 ×g for 10 min, washed twice in PBS, and resuspended in fresh TYG medium without thymidine. The optical density of the cell suspension was adjusted to 16 with the medium, and *B. thetaiotaomicron* and *E. coli* were mixed at equal volumes. Then, 50μL of the mixtures were transferred onto a 0.45 μm filter disc placed on TYG agar plates without thymidine. Plates were incubated anaerobically at 37℃ overnight. To evaluate the number of cells which had acquired plasmids with the transgenes and the erythromycin resistance gene, cells were washed off the filter in the BHIS medium with gentamicin, thymidine, and erythromycin. The cell suspension was plated on the BHIS agar plates containing the same components. The colonies were counted manually 4 days after culturing.

DNA was purified from transconjugants on BHIS agar plates with gentamicin, thymidine, and erythromycin after conjugation, using the DNeasy Blood & Tissue Kits (Qiagen). PCR was performed with primers Seq-*thyA*-F and mmD663 to detect the region including the CD, and qPCR-EmR-F and qPCR-EmR-F for the erythromycin resistance gene (**Supplementary Table 6**). Whole genome sequencing was also performed with the purified DNA to check the integration of the plasmid bearing the transgenes and erythromycin resistance gene.

### Whole genome sequencing and assembly

Total DNA of erythromycin-resistant colonies was purified using the DNeasy Blood & Tissue Kits (Qiagen). Genomic DNA library preparation and sequencing were performed at the Microbial Genome Sequencing Center (MiGS; Pittsburgh, PA) by using Illumina NextSeq 2000 technology. Paired-end sequencing reads (2×151 bp) were assembled by SPAdes 3.15.2^50^. Raw sequencing data and assembled genomes have been deposited at DDBJ/ENA/GenBank under the accession number: PRJNA754595.

### Evolutionary stability test

The stability of the CD and ER during cell growth was verified by culturing the genetically modified thymidine-auxotrophic *B. thetaiotaomicron* strain with those functions up to 21 days. The expression of the NanoLuc gene and the ability to prohibit the HGT of the *thyA* gene were checked every 7 days. Overnight cultures in TYG medium with thymidine were diluted 1:100000 in fresh medium and grown anaerobically at 37℃ for 24 hrs, then grown continuously with dilution into fresh TYG media with thymidine to 10^-5^ of optical density at 600nm every 24±2.5hrs^32^. The functioning of the CD and ER was evaluated on samples withdrawn from the passaged culture in accordance with the protocols of the NanoLuc luciferase assay and the assay for frequency of HGT of the *thyA* gene mentioned above. Samples were grown in continuous culture and measured periodically, as described.

DNA was purified from transconjugants on BHIS agar plates with gentamicin, thymidine, and erythromycin after conjugation, using the DNeasy Blood & Tissue Kits (Qiagen). PCR was performed with primers Seq-*thyA*-F and mmD663 to amplify the region including the CD, and qPCR-EmR-F and qPCR-EmR-F to detect the erythromycin resistance gene (**Supplementary Table 6**). Sanger sequencing of the portion from the P_cepA_-sgRNA1 to the terminator downstream of SpCas9 was performed with the purified amplicons to check if there were any mutations in the CD region. Whole genome sequencing was also performed with the purified DNA to check the integration of the plasmid bearing the intact *thyA* gene.

### Animals

The protocols of all animal experiments were approved by the MIT Committee on Animal Care. Specific-pathogen-free female BALB/cJ mice (8 weeks) were purchased from The Jackson Laboratory, and four mice in each experimental group were housed together and handled in non-sterile conditions. Mice were fed an irradiated mouse diet (ProLab IsoPro RMH3000, LabDiet) and non-sterile water before treatment with antibiotics. Prior to bacterial administration, mice were transferred to clean cages and then gavaged with metronidazole (100mg/kg) in sterile water every day for 7 days. Over the course of the treatment, they were provided sterile filtered water containing ciprofloxacin hydrochloride (0.625 g/L). Animals were transferred to clean cages and provided fresh medicated water on the third or fourth day of the antibiotic regimen. Two days after the cessation of antibiotic treatment, they were transferred to clean cages and inoculated with *B. thetaiotaomicron* strains (∼4×10^8^ CFU/mouse) by oral administration. Fecal pellets were collected for the NanoLuc luciferase assay and the viability test of the bacterial strains in mice feces.

### NanoLuc luciferase assay *in vivo*

Fecal pellets were homogenized in PBS with an autoclaved single 5mm stainless steel bead (Qiagen) using TissueLyser II (Qiagen) at 25Hz for 2mins. The concentration of feces was adjusted to less than 27 mg/mL. The fecal suspensions were centrifuged at 500 ×g for 30s to precipitate debris. The supernatants were diluted 10-fold in PBS. NanoLuc Reaction Buffer (Promega) and supernatants were mixed at equal volumes, and luciferase activities were measured with an integration time of 1 second at a gain setting of 100 in BioTek Synergy H1 Hybrid Reader 6 mins after mixing. The luminescent values were normalized to the weight of the stools.

### Viability test of genetically modified thymidine-auxotrophic *B. thetaiotaomicron in vivo*

Fecal pellets were collected and stored in the presence of oxygen at ambient temperature prior to evaluation. Pellets were homogenized in PBS with a single autoclaved 5mm stainless steel bead (Qiagen) using TissueLyser II (Qiagen) at 25Hz for 2mins. The fecal suspensions were centrifuged at 500 ×g for 30s to precipitate debris.

Supernatants were serially diluted under anaerobic conditions. The dilutions were plated on TYG agar plates with erythromycin, gentamicin, carbenicillin, and polymyxin B, or TYG agar plates with erythromycin, gentamicin, carbenicillin, polymyxin B, and thymidine. Luminescent colonies were counted manually when the NanoLuc Reaction Buffer (Promega) was spotted on the colonies, four days after culturing. The CFU values were normalized to the weight of the stools.

### Statistics

All data were analyzed using GraphPad Prism (GraphPad Software).

## Data availability

Data supporting this study are presented in the main text and Supplementary information, and available from the corresponding author upon request. Plasmids used in this study are available from the corresponding author upon request.

## Acknowledgements

We would like to thank all lab members in the Synthetic Biology Group in MIT’s Synthetic Biology Center for their great help, especially, Dr. Ky Lowenhaupt, Dr. Isaak E. Mueller, Dr. Giyoung Jung, Dr. Tzu-Chieh Tang, Dr. Kevin M. Yehl and Dr. Maria Eugenia Inda. We thank Karen Pepper for editing the manuscript. We thank Dr. Xiaoqiong Gu for bioinformatics analysis. We also thank the staff at the Koch Institute Swanson Biotechnology Center and the Division of Comparative Medicine at MIT for their assistance for animal experiments. This study was supported by JSR Corporation in Japan. This work was supported by the National Science Foundation (NSF-CCF-1521925 to T.K.L.) and the National Institutes of Health (NIH-5-U01-CA2550554-02 and NIH-50000655-5500001351 to T.K.L.).

## Author contributions

N.H., Y. L, M. M. and T. K. L. conceived and designed this study. N. H. and Y. L. performed the experiments. N.H., Y. L., M. M. and T. K. L. analyzed data, discussed the results, and wrote the manuscript.

## DECLARATION OF INTERESTS

T.K.L. is a co-founder of Senti Biosciences, Synlogic, Engine Biosciences, Tango Therapeutics, Corvium, BiomX, Eligo Biosciences, Bota.Bio, and Avendesora. T.K.L. also holds financial interests in nest.bio, Ampliphi, IndieBio, MedicusTek, Quark Biosciences, Personal Genomics, Thryve, Lexent Bio, MitoLab, Vulcan, Serotiny, and Avendesora. N.H is an employee of JSR Corporation. Other authors declare no competing interests.

---

**Supplementary Fig. 1.**
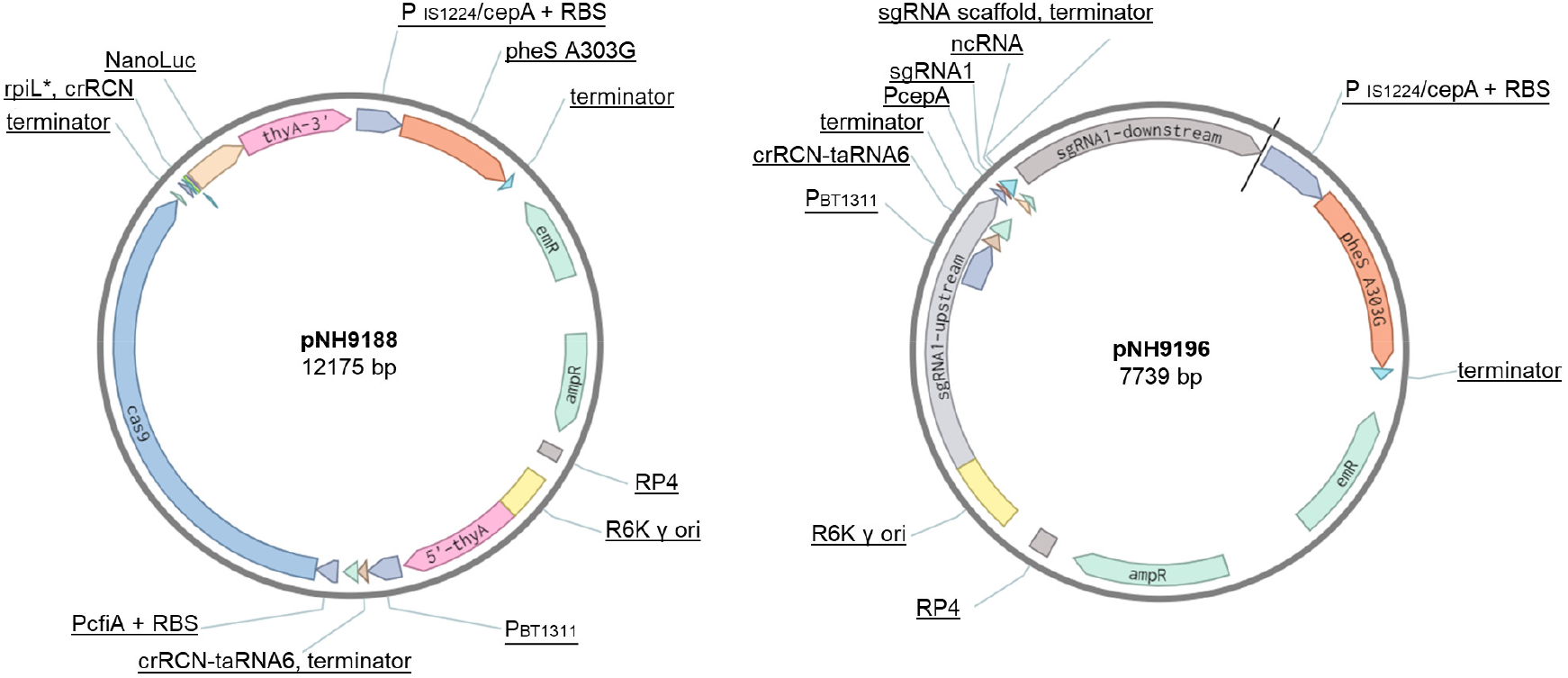
Genetic maps of representative plasmids used in this study. Left, pNH9188, used for the first recombination step, to integrate the components of the Engineered Riboregulator, i.e., the trans-activating RNA sequence (taRNA6) and the cis-repressive sequence (crRCN), and the SpCas9 gene, the NanoLuc gene, and their promoters. Right, pNH9196, used for the second recombination step, to integrate the single-guide RNA sequence (sgRNA1) and its promoter (P_cepA_).

**Supplementary Fig. 2.**
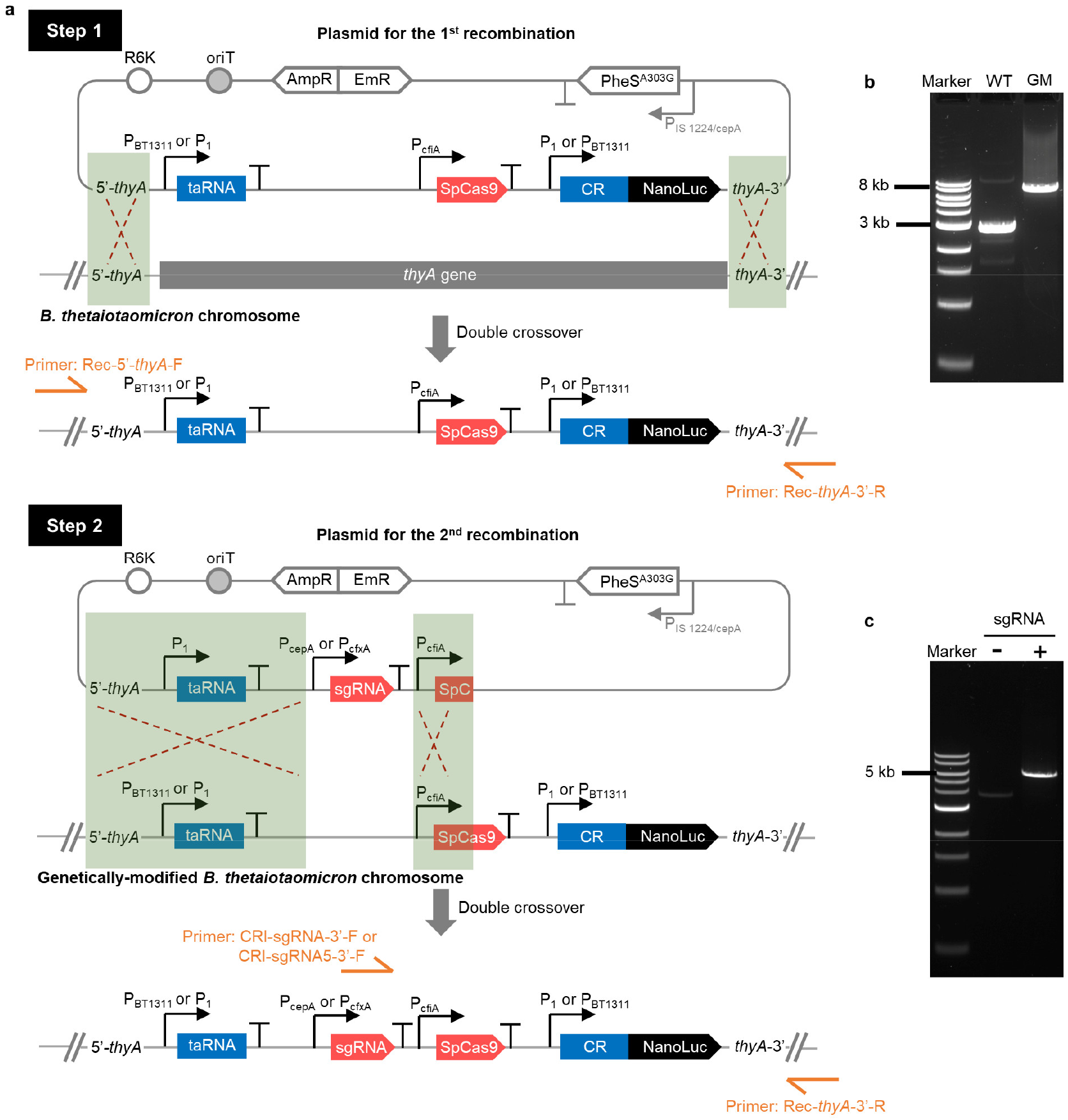
Strategy of genetic modification. **a**, Overview of the recombination method used in this study. Step 1: Introduction of the taRNA, SpCas9, cis-repressive sequence (CR), and NanoLuc gene onto the chromosome of wild-type (WT) *B. thetaiotaomicron*. Step 2: Introduction of sgRNA and its promoters. R6K, origin of replication; oriT, origin of transfer; AmpR, ampicillin resistance cassette; EmR, erythromycin resistance cassette; PheS^A303G^, mutated α−subunit of phenylalanyl-tRNA synthetase gene. **b**, PCR identification of the integration of the gene cassette with primers shown in (**a**). **c**, PCR identification of the integration of sgRNA.

**Supplementary Fig. 3.**
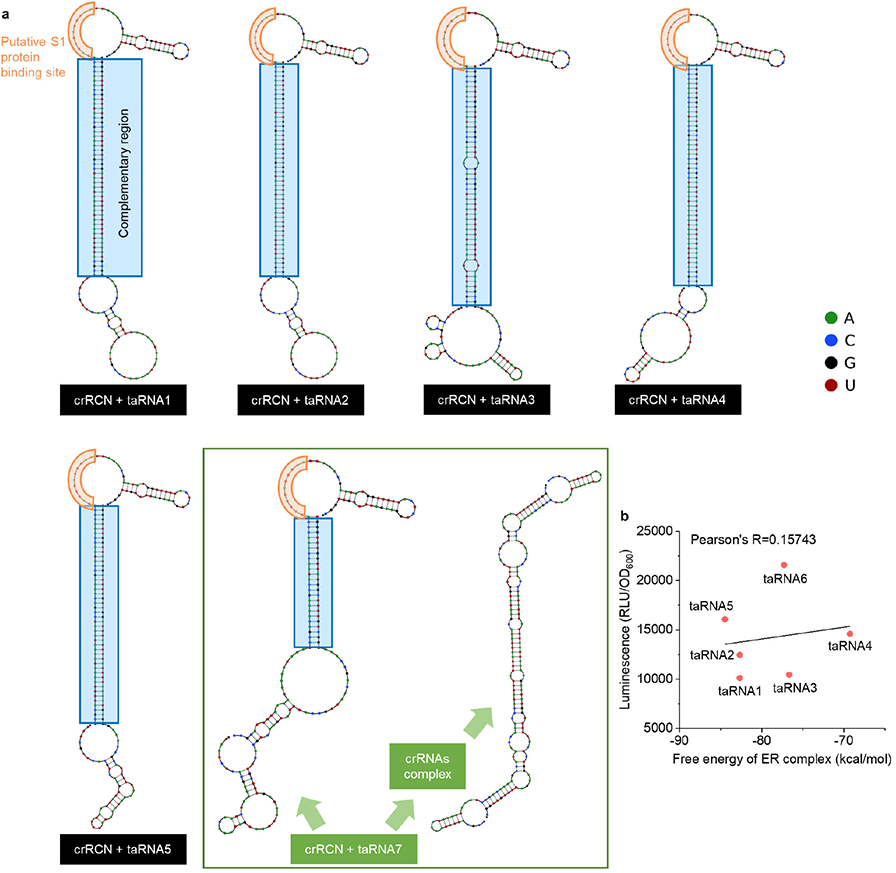
Analysis of crRNA (crRCN)/ taRNA complexes with NUPACK. **a**, NUPACK-predicted structures of crRNA (crRCN)/ taRNA complexes. **b**, Correlation between Minimum Free Energy (i.e., free energy of ER complex) and bioluminescent signal of each crRNA (crRCN)/ taRNA complex: crRNA (crRCN)/ taRNA1 to crRNA (crRCN)/ taRNA6.

**Supplementary Fig. 4.**
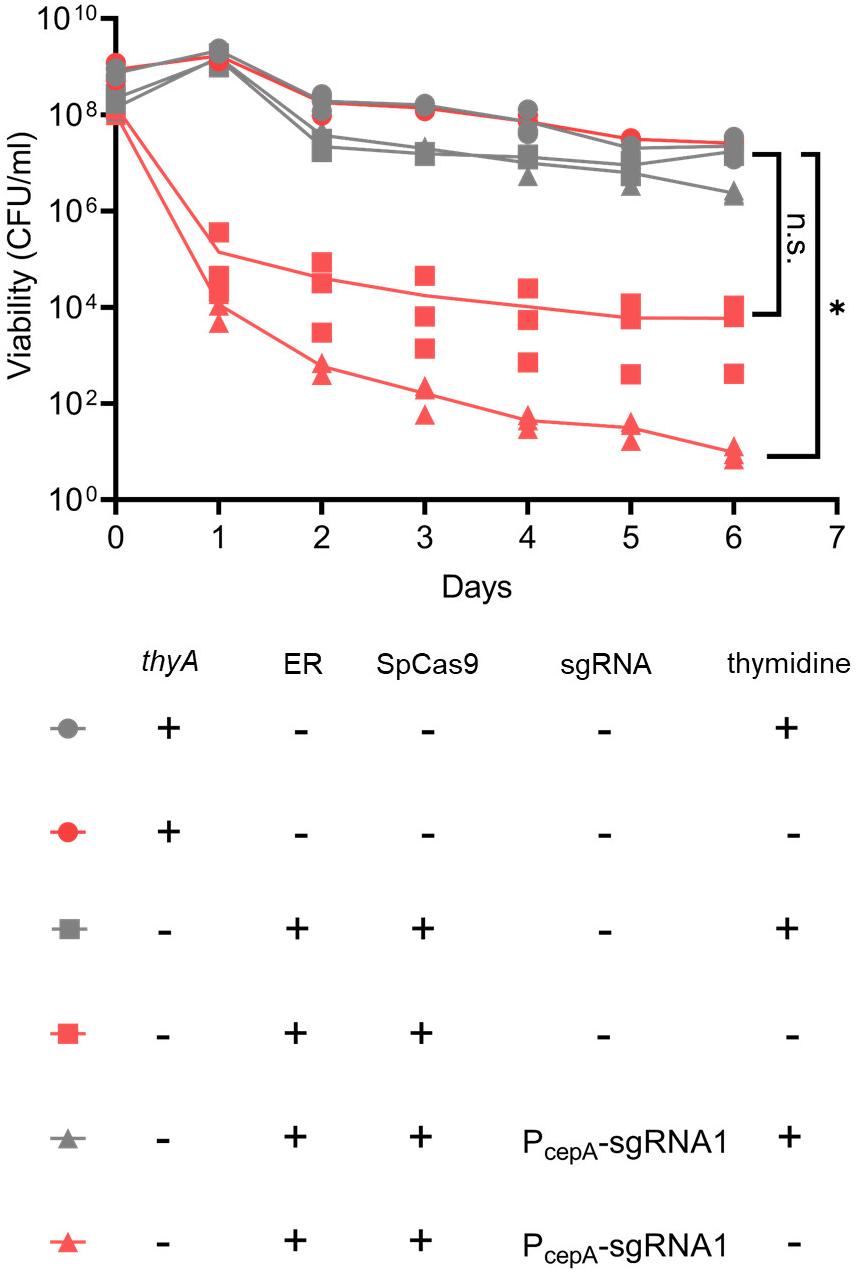
Death rate accelerated by CRISPR Device. Cell viability of WT strain and genetically modified strain with and without P_cepA_-sgRNA1, cultured anaerobically in the presence and absence of thymidine. Each dot is a biological replicate, and lines are the values of the mean of three (Dunn’s multiple comparisons test with Kruskal-Wallis test on log-transformed data at day6, *P<0.05). n.s. represents not significant.

**Supplementary Fig. 5.**
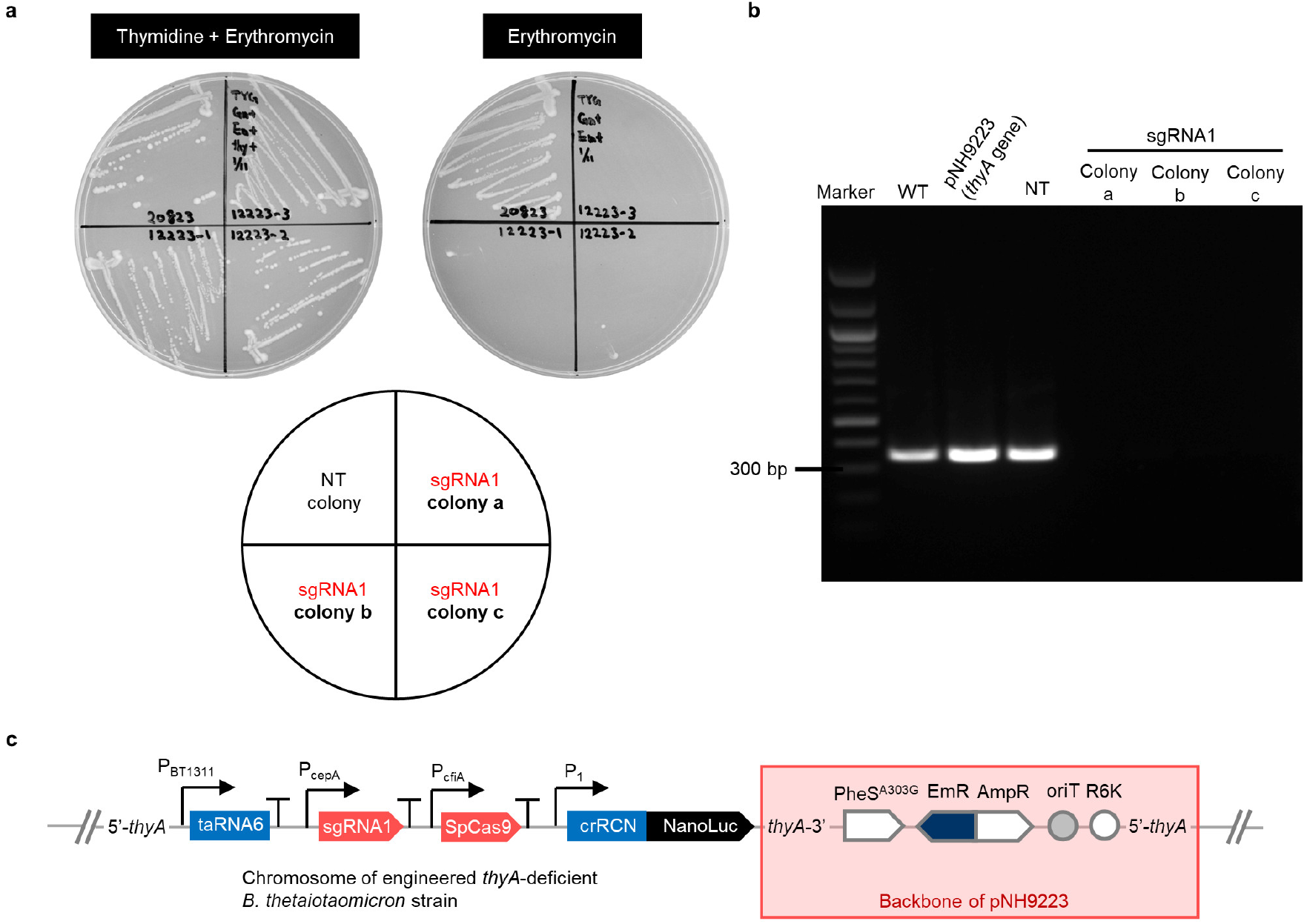
Prevention of acquisition of *thyA* gene by genetically engineered strain. **a**, Colonies streaked on TYG with and without thymidine in the presence of erythromycin and gentamicin. Viable *B. thetaiotaomicron* bearing sgRNA1 or NT, plated on BHIS agar plates with thymidine, erythromycin and gentamicin after conjugation, were streaked to check thymidine auxotrophy. **b**, Amplicon patterns of transconjugants isolated from BHIS agar plates with thymidine, erythromycin, and gentamicin after conjugation. PCR was performed with primers flanking the sequence targeted by the CRISPR Device with sgRNA1 in order to detect the *thyA* gene. **c**, Integration of pNH9223 backbone into the chromosome of engineered *thyA*-deficient *B. thetaiotaomicron* strain.

**Supplementary Fig. 6.**
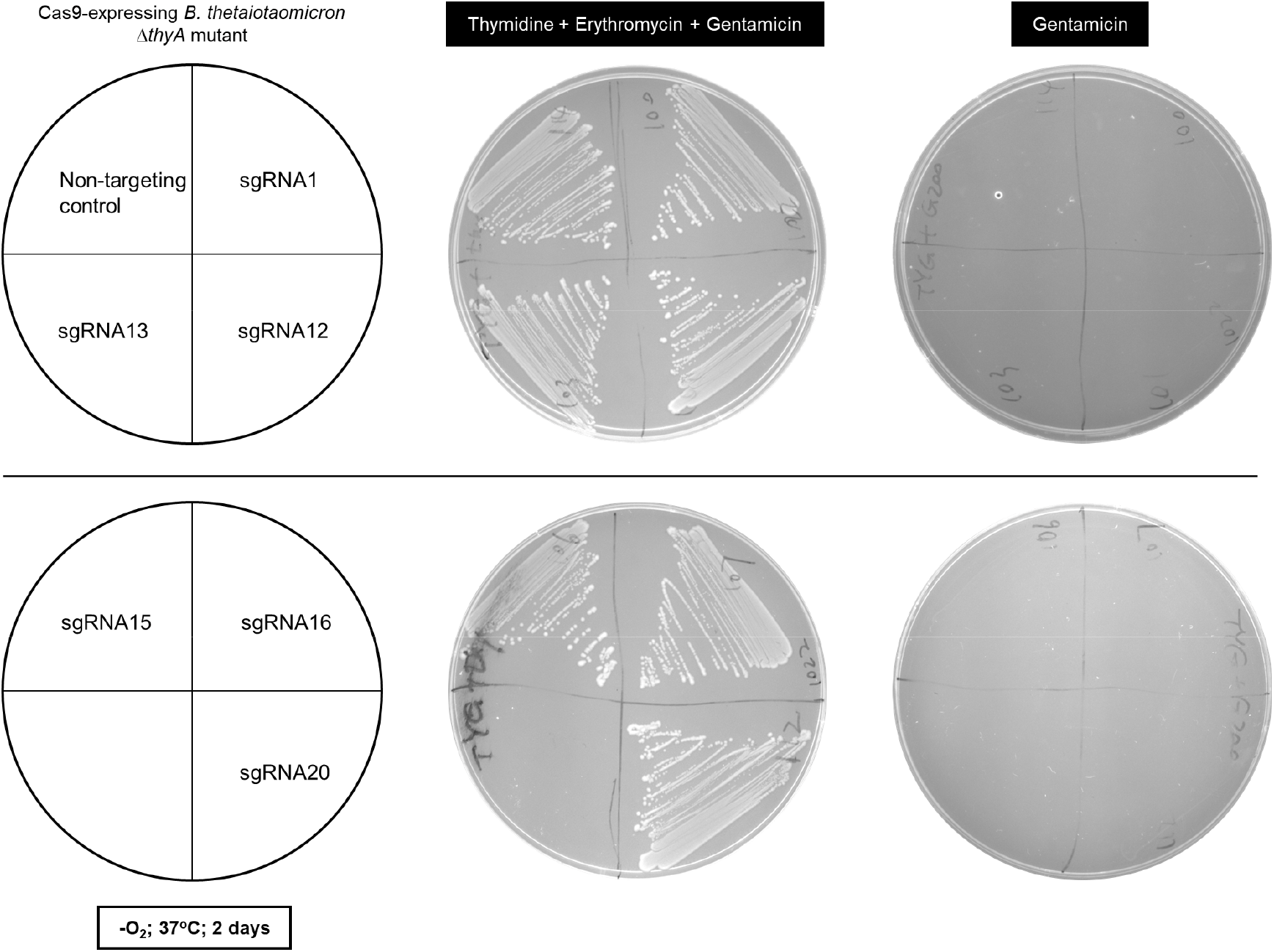
Thymidine auxotrophy of genetically engineered strains bearing the sgRNAs and their promoter constructed by pNBU2-based technology after the first recombination. The genetically engineered strains were streaked on gentamicin-containing TYG agar with both thymidine and erythromycin, or without either thymidine or erythromycin. Plates were anaerobically incubated at 37°C for 2 days.

**Supplementary Fig. 7.**
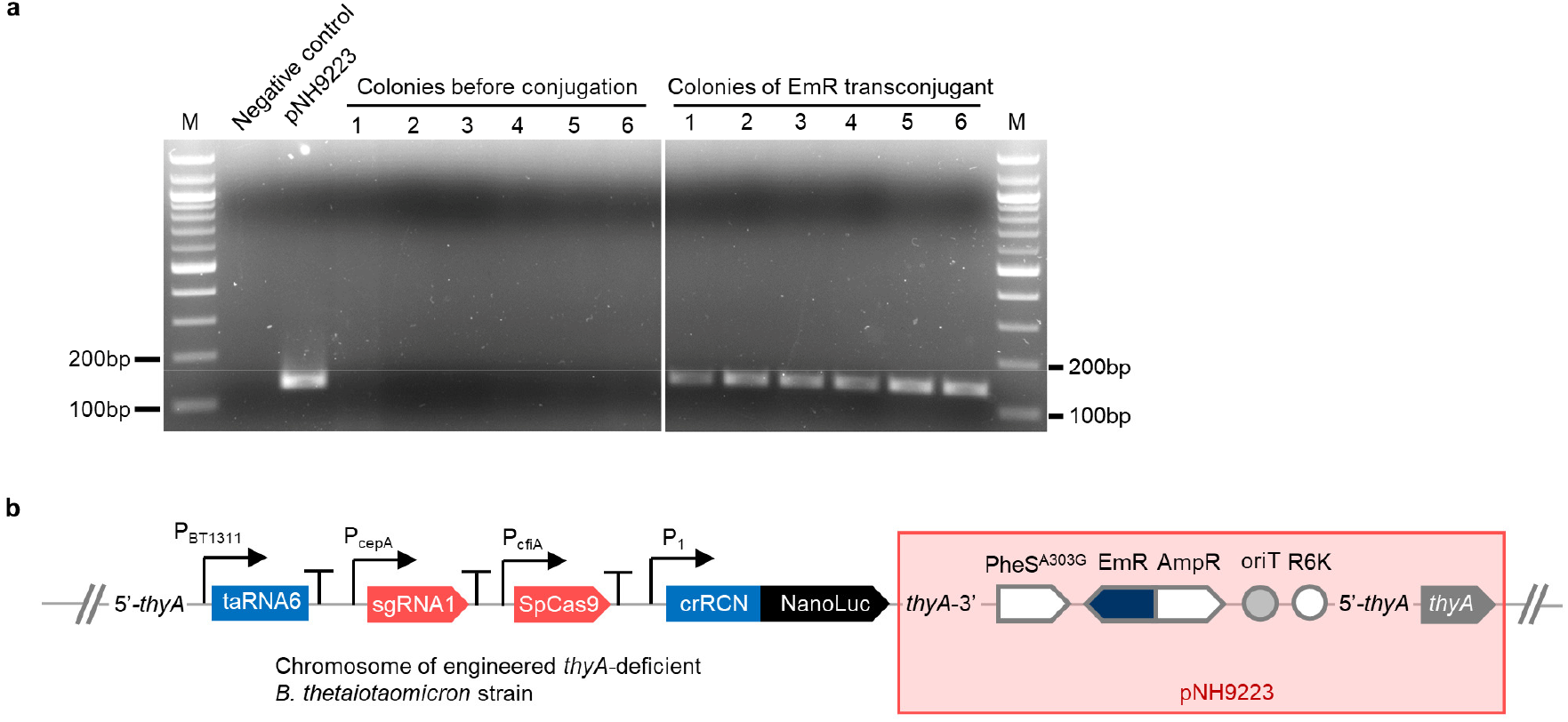
Integration of EmR and *thyA* genes in engineered *thyA*-deficient *B. thetaiotaomicron* strain after conjugation at day21. **a**, PCR amplification of EmR gene fragment (144 bp) in engineered *B. thetaiotaomicron* strains before and after conjugation of pNH9223. **b**, Integration of pNH9223 plasmid into the chromosome of engineered *thyA*-deficient *B. thetaiotaomicron* strain.

**Supplementary Fig. 8.**
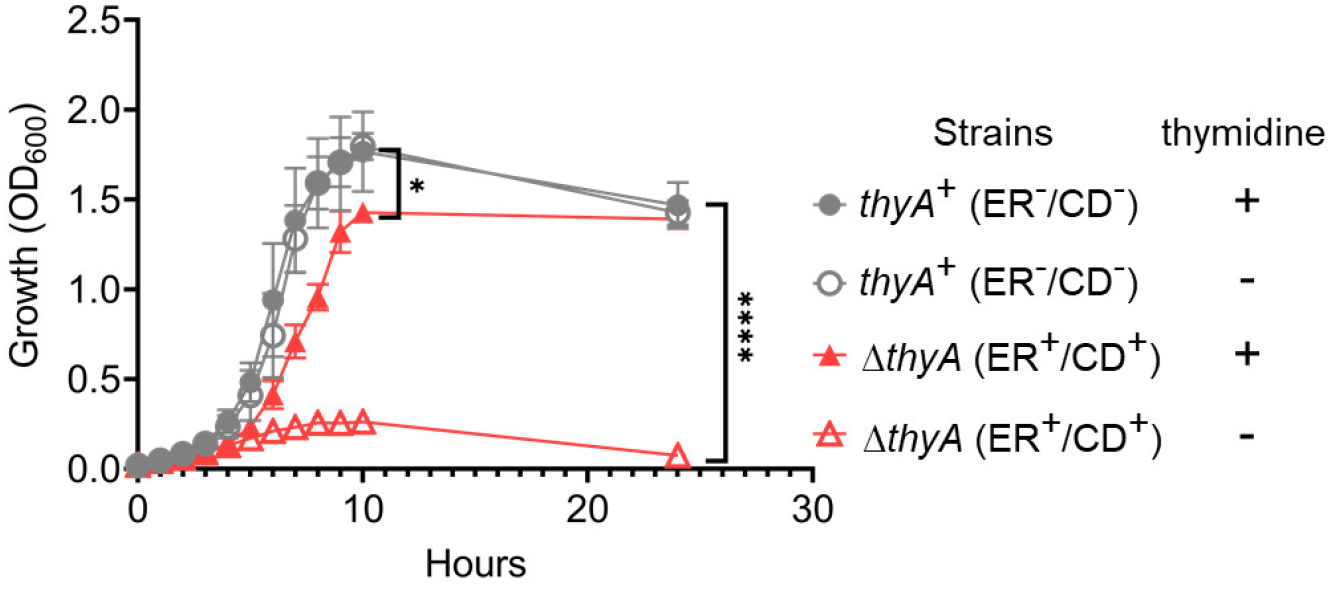
Growth curves of *thyA*^+^ (ER^-^/CD^-^) and Δ*thyA* (ER^+^/CD^+^) strains in the presence and absence of thymidine. The cells were grown anaerobically at 37℃ for 24 hrs. 300 μl of samples were withdrawn from the cultures every hour for 10hrs and at 24hrs to monitor growth (OD_600_). Each dot is the value of the mean, and error bars represent the standard deviations of three biological replicates made on different days (Tukey’s multiple comparisons test with one-way analysis of variance on data at 10hrs and 24hrs, * P<0.05, **** P<0.0001).

**Supplementary Fig. 9.**
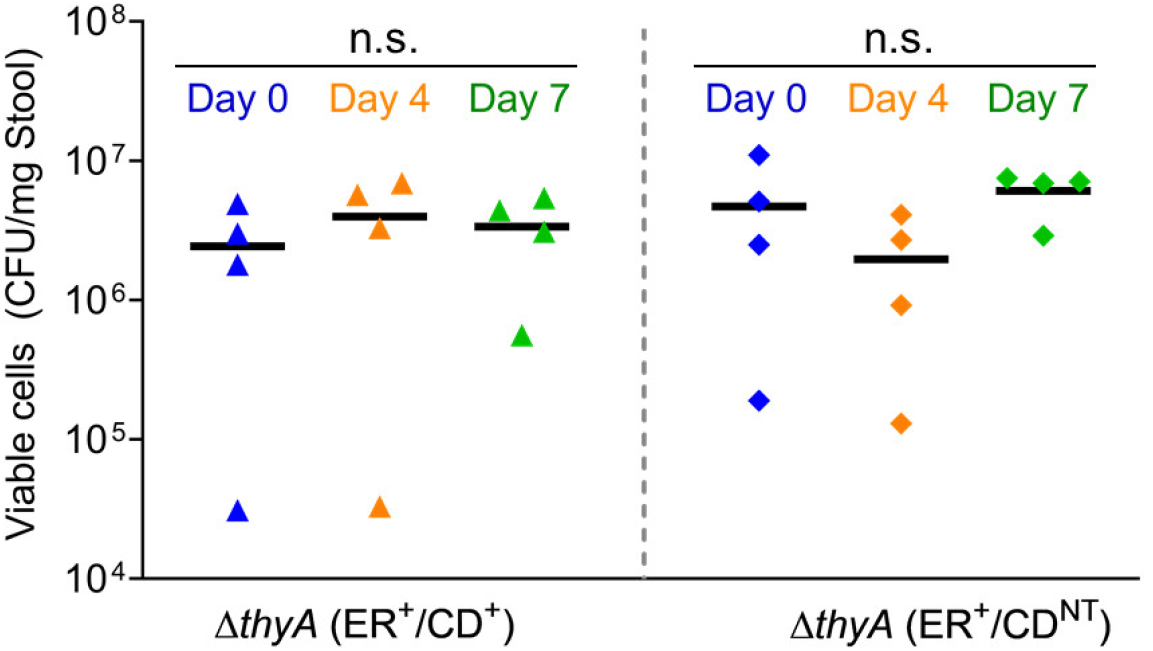
Viability of genetically engineered thymidine-auxotrophic *B. thetaiotaomicron* with engineered riboregulator and CRISPR device in mice feces. Fecal suspensions from mice gavaged with either of Δ*thyA* (ER^+^/CD^+^) or Δ*thyA* (ER^+^/CD^NT^) strains without *thyA*^+^ (ER^-^/CD^-^) strain were plated on selective agar plates with thymidine on the day when feces were retrieved, and after four and seven days of storage at ambient temperature in the presence of oxygen. Lines are the values of the mean of four biological replicates (Tukey’s multiple comparisons test with one-way analysis of variance on log-transformed data). n.s. represents not significant.

