## Supplementary Table 1 for "Cas9-Assisted Biological Containment of a Genetically Engineered Human Commensal Bacterium and Genetic Elements"

**Supplementary Table 1. List of CRs used in Figure 2.**

| **Name** | **Sequence** |
|---|---|
| crRCN | ccgacaaaagccataaccaaaccca |
| crR7N | tcgtttaattaaattcatattgggt |
| crR10N | tcgttaaataaaatacatattgggt |
| crR12N | tcgtataaataaatacatattgggt |
