## Supplementary Table 2 for "Cas9-Assisted Biological Containment of a Genetically Engineered Human Commensal Bacterium and Genetic Elements"

**Supplementary Table 2. List of taRNAs used in Figure 2.**

| Name | Description | Sequence |
|---|---|---|
| Non Sense (NS) | Non-sense trans-activating RNA | taatacccaaatccaggaggtgattggtagtggtggttaatgaaaattaacttactactaccatatatctctaga |
| taRNA1 | trans-activating RNA | tgggtttggttatggcttttgtcggatttattttattttaaaatgcgggaaaccgcatttttaaaataaataaaatccgtaa |
| taRNA2 | trans-activating RNA | tgggtttggttatggcttttgtcggatttattttattttaaaatgcgggaaaccgcatttttaaaataaatattttccgtaa |
| taRNA3 | trans-activating RNA | tgggtttggttatggctttagtcggatttattttattttaaaatgcgggaaaccgcatttataaaataaataaaatccgtaa |
| taRNA4 | trans-activating RNA | tgggtttggttatggcttttgtcggatttattttattttaaaatgcgggaaaccgcatttttaaaataaatattttccgtat |
| taRNA5 | trans-activating RNA | tgggtttggttatggcttttgtcggatttattttattttaaaatgcgggaaaccgcatttttaaaataaatattttaattat |
| taRNA6 | trans-activating RNA | tgggtttggttatggctttagtcggatttattttattttaaaatgcgggaaaccgcatttataaaataaatattttaattat |
| taRNA7 | trans-activating RNA | tgggtttggttatggcttttgtcgggaaaccgacaattgccattta |

Complementary sequences to crRNA are highlighted in blue.
