## Supplementary Table 3 for "Cas9-Assisted Biological Containment of a Genetically Engineered Human Commensal Bacterium and Genetic Elements"

**Supplementary Table 3. List of sgRNAs used in the experiments.**

| Name | Description | Figure | Sequence | PAM |
|---|---|---|---|---|
| Non-targeting control (NT) | Non-targeting single guide RNA | 4, 5, 7, S5, S6, S9 | attctctagactttacttga | None |
| sgRNA1 | Single guide RNA | 3, 4, 5, 6, 7, S1, S4, S5, S6, S7, S8, S9 | acgcatctggaacgaatggg | CGG |
| sgRNA12 | Single guide RNA | 4, S6 | aggcgctgacaatgatacgg | CGG |
| sgRNA13 | Single guide RNA | 4, S6 | atgcgtttcaacctcgacga | GGG |
| sgRNA15 | Single guide RNA | 4, S6 | actgagaaaagtgaccgtac | AGG |
| sgRNA16 | Single guide RNA | 4, S6 | agttttacgtggcagacggt | CGG |
| sgRNA20 | Single guide RNA | 4, S6 | gcctcaaatgaaaattaatc | CGG |
