## Supplementary Table 4 for "Cas9-Assisted Biological Containment of a Genetically Engineered Human Commensal Bacterium and Genetic Elements"

**Supplementary Table 4. List of genetic parts used in experiments.**

| Name | Description (Source) | Sequence |
|---|---|---|
| AmpR | Ampicillin-resistance cassette (Cell Syst. 2015; 1(1): 62-71.) | atgagtattcaacatttccgtgtcgcccttattcccttttttgcggcattttgccttcctgtttttgctcacccagaaacgctggtgaaagtaaaagatgctgaagatcagttgggtgcacgagtgggttacatcgaactggatctcaacagcggtaagatccttgagagttttcgccccgaagaacgttttccaatgatgagcacttttaaagttctgctatgtggcgcggtattatcccgtattgacgccgggcaagagcaactcggtcgccgcatacactattctcagaatgacttggttgagtactcaccagtcacagaaaagcatcttacggatggcatgacagtaagagaattatgcagtgctgccataaccatgagtgataacactgcggccaacttacttctgacaacgatcggaggaccgaaggagctaaccgcttttttgcacaacatgggggatcatgtaactcgccttgatcgttgggaaccggagctgaatgaagccataccaaacgacgagcgtgacaccacgatgcctgtagcaatggcaacaacgttgcgcaaactattaactggcgaactacttactctagcttcccggcaacaattaatagactggatggaggcggataaagttgcaggaccacttctgcgctcggcccttccggctggctggtttattgctgataaatctggagccggtgagcgtgggtctcgcggtatcattgcagcactggggccagatggtaagccctcccgtatcgtagttatctacacgacggggagtcaggcaactatggatgaacgaaatagacagatcgctgagataggtgcctcactgattaagcattggtaa |
| EmR | Erythromycin-resistance cassette (Cell Syst. 2015; 1(1): 62-71) | atgaacaaagtaaatataaaagatagtcaaaattttattacttcaaaatatcacatagaaaaaataatgaattgcataagtttagatgaaaaagataacatctttgaaataggtgcagggaaaggtcattttactgctggattggtaaagagatgtaattttgtaacggcgatagaaattgattctaaattatgtgaggtaactcgtaataagctcttaaattatcctaactatcaaatagtaaatgatgatatactgaaatttacatttcctagccacaatccatataaaatatttggcagcataccttacaacataagcacaaatataattcgaaaaattgtttttgaaagttcagccacaataagttatttaatagtggaatatggttttgctaaaatgttattagatacaaacagatcactagcattgctgttaatggcagaggtagatatttctatattagcaaaaattcctaggtattatttccatccaaaacctaaagtggatagcacattaattgtattaaaaagaaagccagcaaaaatggcatttaaagagagaaaaaaatatgaaacttttgtaatgaaatgggttaacaaagagtacgaaaaactgtttacaaaaaatcaatttaataaagctttaaaacatgcgagaatatatgatataaacaatattagtttcgaacaatttgtatcgctatttaatagttataaaatatttaacggctaa |
| R6K | Origin of replication (Cell Syst. 2015; 1(1): 62-71) | gatctgaagatcagcagttcaacctgttgatagtacgtactaagctctcatgtttcacgtactaagctctcatgtttaacgtactaagctctcatgtttaacgaactaaaccctcatggctaacgtactaagctctcatggctaacgtactaagctctcatgtttcacgtactaagctctcatgtttgaacaataaaattaatataaatcagcaacttaaatagcctctaaggttttaagttttataagaaaaaaaagaatatataaggcttttaaagcttttaaggtttaacggttgtggacaacaagccagggatgtaacgcactgagaagcccttagagcctctcaaagcaattttgagtgacacaggaacacttaacggctgaca |
| RP4 | Origin of transfer (Cell Syst. 2015; 1(1): 62-71) | ccggccagcctcgcagagcaggattcccgttgagcaccgccaggtgcgaataagggacagtgaagaaggaacacccgctcgcgggtgggcctacttcacctatcctgccc |

| | | |
|---|---|---|
| PheS$^{A}_{303G}$ | Mutated alpha subunit of Bacteroides phenylalanyl tRNA synthetase (Anaerobe. 2016; 42: 81-88.) | atgatagctaagattaatcaacttcttgaagaggtgggggccttgaaagcc<br>gccaatgccgaagaactcgaagttctgcgcatcaaatacctcagcaaga<br>aaggagccatcaatgacttgatggcagatttccgcaatgtggctgccgaa<br>cagaaaaaagaagtcggcatgaagctcaacgagctgaaaacaaaag<br>cacaagaaaaaatcaacgcactgaaagaacagttcgacaaccaggac<br>aacggacaggacgatctcgacctgacccgttcggcttatcccgtggagct<br>gggcacacgccatccgctttccattgtgcgcaacgaaatcattgacatcttt<br>gcccgtctgggattcaacattgccgaaggtccggaaatagaagatgact<br>ggcatgtgttctccgcactcaactttgccgaagaccatccggcacgagac<br>atgcaagacacattcttcatcgaatcccatcccgacgtactactgcgcaca<br>cacacctcatcggtacagagccgtgtgatggaggtttcgcaaccacctat<br>ccgtatcatctgtccgggacgcgtttaccgtaacgaagctatcagctatcgt<br>gcacactgtttcttccaccaggtagaagcgctgtacgtagaccgcaatgta<br>tctttcaccgacctgaaacaagtgttgctactcttcgccaaagagatgttcg<br>gtgccgatacgaagattcgcctgcgtccgtcgtacttcccgttcacagaac<br>ccagcgccgaaatggatatcagctgtaacatctgcggcggaaaaggctg<br>ccccttctgcaaacacaccggctgggtcgaaatcttgggttgcggaatggt<br>agacccgaacgtacttgatgccaacggaatagacagcaaagtgtatag<br>cggatatggactgggtatgggtatcgaacgcatcacaaacctgaaatatc<br>aggtgaaagacctccgcatgttctccgaaaacgatacacgtttcctgaag<br>gagttcgaagcggcctattaa |
| $P_{IS\ 1224/cepA}$ + RBS | Constitutive promoter + RBS (Anaerobe. 2016; 42: 81-88.) | cctttgaaaagagaagaagccgtgttgctgttcaaactggtcaatgacttc<br>caggaaaggacatcactcatcatcacggcaaacaaggcactcacccgt<br>tggctggaaacattggaggatgaagcggtcacagccgccttgcttgaca<br>ggctgctctactgctgcgagattatcaggctcggaggaacaagctatcgc<br>atgcaaaacaggaaaacaatttttagcaaccaaaacacggatataggc<br>acgtaaaaagagttaaggaaagtgaagcatcttcgatgctggagtggga<br>tatactaaattacctaaaaaagtggcgcacaaatttgcgcgccacaattat<br>tattcatacctttgtggaccgtattacaaagaacccaatcatat |
| $P_{BT1311}$ | Constitutive promoter (Cell Syst. 2015; 1(1): 62-71) | tgatctggaagaagcaatgaaagctgctgttaagtctccgaatcaggtatt<br>gttcctgacaggtgtattcccatccggtaaacgcggatactttgcagttgatc<br>tgactcaggaataaattataaattaaggtaagaagattgtaggataagcta<br>atgaaatagaaaaaggatgccgtcacacaacttgtcggcattctttttgtttt<br>attagttgaaaatatagtgaaaaagttgcctaaatatgtatgttaacaaatta<br>tttgtcgtaactttgcactc |
| $P_1$ | Constitutive promoter (Cell Syst. 2015; 1(1): 62-71) | gataaagtttggaagataaagctaaaagttcttatctttgcagt |
| $P_{cfiA}$ + RBS | Constitutive promoter + RBS (Cell Syst. 2015; 1(1): 62-71) | ggagtgagcttctcggattttatttgtatttttgccatgcctgatgaggttttgtttg<br>attattttttgcaacactaagttaagtgaatcctctgacatggcaaaatcctg<br>agcaactttttgttgctcaggtacttaaaaaaaatattttataatagtgttgcg<br>gaattaaggtaaaagaataaa |
| $P_{cfxA}$ | Constitutive promoter (Cell Syst. 2015; 1(1): 62-71) | tacaaagaaaattcgacaaactgttatttttctatctatttatttgggtgggaa<br>actttagttatgtacctttgtcggc |
| $P_{cepA}$ | Constitutive promoter (Cell Syst. 2015; 1(1): 62-71) | caaatttgcgcgccacaattattattcatacctttgtgg |
| rpiL* | RBS (Cell Syst. 2015; 1(1): 62-71) | ccgcattttaaaataaaataaattatttatttaattaaacgaat |

| | | |
|---|---|---|
| 5'-*thyA* | Upstream of *thyA* gene of *B. thetaiotaomicron* (This work) | tctatagccaatgcgctgttcgacctgctggccgaaaaagtgaaagaaggcgtggaagtacgagccatgttcgatgcgttcggcaactggtcgaacaataaaccgctcaagaagagacatctcaaaaaaatacgagagcagggaatcgagattgtcaagttcgacccgttcacctttccctacatcaatcatgcggcacaccgtgatcaccgcaaaatagccgtcatcgacggtgaagtcgcctatacgggaggaatgaacatcgccgactactacatcaacggactccccaaaatcggtacatggcgtgatatgcacatgagaatagaaggcgatgccgtcaatgacctgcaagaaatatttcttaccatatggaacaaggagactaaacaaaatattggcggcgaagcatacttccccaagcacaaggagcagtcggacagcaccaatgtagtcgtcgccatcgtagaccgcacaccgaagaaaaacagccgtatgctcagtcatgcttatgccatgtcgatctattcggcacagaaaaacgtgcacatcgtcaacccgtattttgtgcccacatcctccatcaacaaggcgcttcaacgcaccatcgaacgaggtgtggatgtcacgattatggtttcttctgcctccgacatcccgtttactccggacgccgccctctacaaacttcacaaactgatgaaaagggggccaccgtctatatgtacaatggcggtttccaccattccaaaatcatgatggtcgacgatattttctgcacggtaggcaccgcaaacctgaacagccgcagcctccggtacgactacgagacgaacgctttcatctttaacaaggaaattacaggcgaactgaacgaaatgtttcggaacgatatagaacactgtacgcagctaacaccggaattctggaaaaagcgctcaccgtggaagaagttcgtcggatggtttgccaatttattcacgccatttttgtaattttgcatcgcaccaacttattggaaaa |
| *thyA*-3' | Downstream of *thyA* gene of B. thetaiotaomicron (This work) | ccacatattgccggaatagtagcggtataaaaccatgagtaaaatatcaattattgccgccgtagaccgccgaatggcgatcggctttcagaataaactgcttttctggttgcccaacgacttgaaacgcttcaaggcactgactaccggaaacaccatcataatgggaagaaaaaccttcgaatcgctgccgaaaggtgcgttgcccaatcgcagaaacgttgtgttatccacccgtccggatacggtatgtcccggtgcggaagtcttcccgtcgctggaagtcgccctgcaaagctgcaaggaagatgagcacgtctatataataggtggcgcaagcgtctaccagcaggcacttcctcttgccgacgaactttgtctgacggaaataaatgacgttgctcccgaagccgatgccttctttccggaagtatctccggctcaatggcacgaaaaaagcagagaagctcatcctgtggatgagaaacatctctgcccgtatgcttttgtagattacgtgaaacagtaagctattttattctgtacgattaatcttcgtcttcaatgtcgaccataatcggcatctgccgctttatggcttcatggaaagagatcaatgtctccgtacgcgccaaccccaaaggttgcaatttatcatgtatgatactcagcaaatggtgattgttcttagcgtaaatcttgataaacatatcatatttaccagtagtgaaatgacactctactacttccggaatggcttccaaagctctcgttacagaatcaaaggattccggatctttcagatatataccaatataagcacaagtttcatatcctatcttctcaggatcgatcacatattccgatcctttcaatattcccagattagtcaacttctgaatgcgctggtggatagccgccccggaaacgttacacgctcttgctacttccaaa |
| CR | Cis-Represive sequence (This work) | See table S1 |
| taRNA | trans-activating RNA (This work) | See table S2 |

| | | |
|---|---|---|
| SpCas9 | Cas9 gene of *Streptococcus pyogenes* modified to work in *B. thetaiotaomicron* (This work) | atggataagaaatactcaataggcttagatatcggcacaaatagcgtcgg<br>atgggcggtgatcactgatgaatataaggttccgtctaaaaagttcaaggtt<br>ctgggaaatacagaccgccacagtatcaaaaaaaatcttataggggctc<br>ttttatttgacagtggagagacagcggaagcgactcgtctcaaacggaca<br>gctcgtagaaggtatacacgtcggaagaatcgtatttgttatctacaggag<br>atttttcaaatgagatggcgaaagtagatgatagtttctttcatcgacttgaa<br>gagtctttttggtggaagaagacaagaagcatgaacgtcatcctatttttgg<br>aaatatagtagatgaagttgcttatcatgagaaatatccaactatctatcatc<br>tgcgaaaaaaattggtagattctactgataaagcggatttgcgcttaatctat<br>ttggccttagcgcatatgattaagtttcgtggtcatttttgattgagggagattt<br>aaatcctgataatagtgatgtggacaaactatttatccagttggtacaaacc<br>tacaatcaattatttgaagaaaaccctattaacgcaagtggagtagatgct<br>aaagcgattctttctgcacgattgagtaaatcaagacgattagaaaatctc<br>attgctcagctccccggtgagaagaaaaatggcttatttgggaatctcattg<br>ctttgtcattgggtttgacccctaattttaaatcaaattttgatttggcagaagat<br>gctaaattacagctttcaaaagatacttacgatgatgatttagataatttattg<br>gcgcaaattggagatcaatatgctgatttgtttttggcagctaagaatttatca<br>gatgctattttactttcagatatcctaagagtaaatactgaaataactaaggc<br>tcccctatcagcttcaatgattaaacgctacgatgaacatcatcaagacttg<br>actcttttaaaagctttagttcgacaacaacttccagaaaagtataaagaa<br>atctttttgatcaatcaaaaaacggatatgcaggttatattgatgggggag<br>ctagccaagaagaattttataaatttatcaaaccaattttagaaaaaatgg<br>atggtactgaggaattattggtgaaactaaatcgtgaagatttgctgcgca<br>agcaacggacctttgacaacggctctattccccatcaaattcacttgggtg<br>agctgcatgctattttgagaagacaagaagacttttatccatttttaaaagac<br>aatcgtgagaagattgaaaaaatcttgacttttcgaattccttattatgttggtc<br>cattggcgcgtggcaatagtcgttttgcatggatgactcggaagtctgaag<br>aaacaattaccccatggaattttgaagaagttgtcgataaaggtgcttcag<br>ctcaatcatttattgaacgcatgacaaactttgataaaaatcttccaaatga<br>aaaagtactaccaaaacatagtttgctttatgagtattttacggtttataacga<br>attgacaaaggtcaaatatgttactgaaggaatgcgaaaaccagcatttct<br>ttcaggtgaacagaagaaagccattgttgatttactcttcaaaacaaatcg<br>aaaagtaaccgttaagcaattaaaagaagattatttcaaaaaaatagaat<br>gttttgatagtgttgaaatttcaggagttgaagatagatttaatgcttcattagg<br>tacctaccatgatttgctaaaaattattaaagataaagatttttggataatga<br>agaaaatgaagatatcttagaggatattgttttaacattgaccttatttgaag<br>atagggagatgattgaggaaagacttaaaacatatgctcacctctttgatg<br>ataaggtgatgaaacagcttaaacgtcgccgttatactggttggggacgttt<br>gtctcgaaaattgattaatggtattagggataagcaatctggcaaaacaat<br>attagatttttgaaatcagatggttttgccaatcgcaattttatgcagctgatc<br>catgatgatagtttgacatttaaagaagacattcaaaaagcacaagtgtct<br>ggacaaggcgatagtttacatgaacatattgcaaatttagctggtagccct<br>gctattaaaaaaggtattttacagactgtaaaagttgttgatgaattggtcaa<br>agtaatggggcggcataagccagaaaatatcgttattgaaatggcacgtg<br>aaaatcagacaactcaaaagggccagaaaaattcgcgagagcgtatg<br>aaacgaatcgaagaaggtatcaaagaattaggaagtcagattcttaaag<br>agcatcctgttgaaaatactcaattgcaaaatgaaaagctctatctctattat<br>ctccaaaatggaagagacatgtatgtggaccaagaattagatattaatcgt<br>ttaagtgattatgatgtcgatcacattgttccacaaagtttccttaaagacgat<br>tcaatagacaataaggtcttaacgcgttctgataaaaatcgtggtaaatcg<br>gataacgttccaagtgaagaagtagtcaaaaagatgaaaaactattgga |

| | | |
|---|---|---|
| | | gacaacttctaaacgccaagttaatcactcaacgtaagtttgataatttaac<br>gaaagctgaacgtggaggtttgagtgaacttgataaagctggttttatcaa<br>acgccaattggttgaaactcgccaaatcactaagcatgtggcacaaatttt<br>ggatagtcgcatgaatactaaatacgatgaaaatgataaacttattcgag<br>aggttaaagtgattaccttaaaatctaaattagtttctgacttccgaaaagatt<br>tccaattctataaagtacgtgagattaacaattaccatcatgcccatgatgc<br>gtatctaaatgccgtcgttggaactgctttgattaagaaatatccaaaacttg<br>aatcggagtttgtctatggtgattataaagtttatgatgttcgtaaaatgattgc<br>taagtctgagcaagaaataggcaaagcaaccgcaaaatatttcttttactc<br>taatatcatgaacttcttcaaaacagaaattacacttgcaaatggagagatt<br>cgcaaacgccctctaatcgaaactaatggggaaactggagaaattgtct<br>gggataaagggcgagattttgccacagtgcgcaaagtattgtccatgccc<br>caagtcaatattgtcaagaaaacagaagtacagacaggcggattctcca<br>aggagtcaattttaccaaaaagaaattcggacaagcttattgctcgtaaaa<br>aagactgggatccaaaaaaatatggtggttttgatagtccaacggtagctt<br>attcagtcctagtggttgctaaggtggaaaaagggaaatcgaagaagtta<br>aaatccgttaaagagttactagggatcacaattatggaaagaagttcctttg<br>aaaaaaatccgattgactttttagaagctaaaggatataaggaagttaaa<br>aaagacttaatcattaaactacctaaatatagtctttttgagttagaaaacgg<br>tcgtaaacggatgctggctagtgccggagaattacaaaaaggaaatgag<br>ctggctctgccaagcaaatatgtgaattttttatatttagctagtcattatgaaa<br>agttgaagggtagtccagaagataacgaacaaaaacaattgtttgtgga<br>gcagcataagcattatttagatgagattattgagcaaatcagtgaattttcta<br>agcgtgttattttagcagatgccaatttagataaagttcttagtgcatataaca<br>aacatagagacaaaccaatacgtgaacaagcagaaaatattattcattta<br>tttacgttgacgaatcttggagctcccgctgcttttaaatattttgatacaaca<br>attgatcgtaaacgatatacgtctacaaaagaagttttagatgccactcttat<br>ccatcaatccatcactggtctttatgaaacacgcattgatttgagtcagctag<br>gaggtgac |
| NanoLuc | NanoLuc gene (Cell Syst. 2015; 1(1): 62-71) | atggtttttactctggaagattttgttggcgattggcgtcagaccgcgggttat<br>aatttggatcaagtcctggaacagggtggcgtaagctctctgttccagaac<br>ctgggtgtgagcgtgacgccgattcagcgcatcgttctgtccggcgagaa<br>cggtctgaaaattgatattcatgtgatcatcccgtacgaaggcctgagcgg<br>tgaccaaatgggtcaaatcgagaaaatctttaaagtcgtctacccagttga<br>cgatcaccacttcaaggttatcttgcattacggtacgctggtgattgatggtg<br>tgaccccgaatatgattgactatttcggccgtccgtatgaaggcattgccgtt<br>tttgacggtaaaaagatcaccgtcaccggtaccctgtggaatggcaataa<br>gattattgacgagcgtctgattaacccggacggcagcctgctgttccgcgt<br>gaccatcaacggtgtcacgggttggcgtctgtgcgagcgcatcctggcat<br>aa |
| sgRNA1-upstream | Upstream of sgRNA (This work) | tctatagccaatgcgctgttcgacctgctggccgaaaaagtgaaagaag<br>gcgtggaagtacgagccatgttcgatgcgttcggcaactggtcgaacaat<br>aaaccgctcaagaagagacatctcaaaaaaatacgagagcagggaat<br>cgagattgtcaagttcgacccgttcacctttccctacatcaatcatgcggca<br>caccgtgatcaccgcaaaatagccgtcatcgacggtgaagtcgcctata<br>cgggaggaatgaacatcgccgactactacatcaacggactccccaaaa<br>tcggtacatggcgtgatatgcacatgagaatagaaggcgatgccgtcaat<br>gacctgcaagaaatatttcttaccatatggaacaaggagactaaacaaa<br>atattggcggcgaagcatacttccccaagcacaaggagcagtcggaca<br>gcaccaatgtagtcgtcgccatcgtagaccgcacaccgaagaaaaaca<br>gccgtatgctcagtcatgcttatgccatgtcgatctattcggcacagaaaaa |

| | | |
|---|---|---|
| | | cgtgcacatcgtcaacccgtattttgtgcccacatcctccatcaacaaggcgcttcaacgcaccatcgaacgaggtgtggatgtcacgattatggtttcttctgcctccgacatcccgtttactccggacgccgccctctacaaacttcacaaactgatgaaaagggggccaccgtctatatgtacaatggcggtttccaccattccaaaatcatgatggtcgacgatattttctgcacggtaggcaccgcaaacctgaacagccgcagcctccggtacgactacgagacgaacgctttcatctttaacaaggaaattacaggcgaactgaacgaaatgtttcggaacgatatagaacactgtacgcagctaacaccggaattctggaaaaagcgctcaccgtggaagaagttcgtcggatggtttgccaatttattcacgccatttttgtaattttgcatcgcaccaacttattggaaaaagcacgtcgtctcagactgaggtcactgagagtagctgatctggaagaagcaatgaaagctgctgttaagtctccgaatcaggtattgttcctgacaggtgtattcccatccggtaaacgcggatactttgcagttgatctgactcaggaataaattataaattaaggtaagaagattgtaggataagctaatgaaatagaaaaaggatgccgtcacacaacttgtcggcattctttttgttttattagttgaaaatatagtgaaaaagttgcctaaatatgtatgttaacaaattatttgtcgtaactttgcactctgggtttggttatggctttagtcggatttattttatttttaaatgcgggaaaccgcatttataaaataaatattttaattattgaactgcacttgctttgataattaatgataaacaatctaaaagcactctaatcgttatcggagtgcttttagattactaatcaaattgcttctactaattgcctatcttccagtgatggaacagcatttgtgcattggctgcaacaacgctaagatgctgg |
| sgRNA1-downstream | Downstream of sgRNA (This work) | ccacgtgctaaggagtgagcttctcggattttatttgtatttttgccatgcctgatgaggttttgtttgattattttttgcaacactaagttaagtgaatcctctgacatggcaaaatcctgagcaactttttgttgctcaggtacttaaaaaaaatattttataatagtgttgcggaattaaggtaaaagaataaaatggataagaaatactcaataggcttagatatcggcacaaatagcgtcggatgggcggtgatcactgatgaatataaggttccgtctaaaaagttcaaggttctgggaaatacagaccgccacagtatcaaaaaaaatcttataggggctcttttatttgacagtggagagacagcggaagcgactcgtctcaaacggacagctcgtagaaggtatacacgtcggaagaatcgtatttgttatctacaggagatttttcaaatgagatggcgaaagtagatgatagtttctttcatcgacttgaagagtctttttggtggaagaagacaagaagcatgaacgtcatcctatttttggaaatatagtagatgaagttgcttatcatgagaaatatccaactatctatcatctgcgaaaaaaattggtagattctactgataaagcggatttgcgcttaatctatttggccttagcgcatatgattaagtttcgtggtcatttttttgattgagggagatttaaatcctgataatagtgatgtggacaaactatttatccagttggtacaaacctacaatcaattatttgaagaaaaccctattaacgcaagtggagtagatgctaaagcgattctttctgcacgattgagtaaatcaagacgattagaaaatctcattgctcagctccccggtgagaagaaaaatggcttatttgggaatctcattgctttgtcattgggtttgacccctaattttaaatcaaattttgatttggcagaagatgctaaattacagctttcaaaagatacttacgatgatgatttagataatttattggcgcaaattggagatcaatatgctgatttgtttttggcagctaagaatttatcagatgctattttactttcagatatcctaagagtaaatactgaaataactaaggctcccctatcagcttcaatgattaaacgctacgatgaacatcatcaagacttgactcttttaaaagctttagttcgacaacaacttccagaaaagtataaagaaatctttttgatcaatcaaaaaacggatatgcaggttatattgatgggggagctagccaagaagaatttttataaatttatcaaaccaattttagaaaaaatggatggtactgaggaattattggtgaaactaaatcgtgaagatttgctgcgcaagc |
| sgRNA | Single guide RNA (This work) | See table S3 |

| | | |
|---|---|---|
| gRNA scaffold | Guide RNA scaffold (Cell Syst. 2015; 1(1): 62-71) | gttttagagctagaaatagcaagttaaaataaggctagtccgttatcaacttgaaaaagtggcaccgagtcggtgc |
| *thyA* | *thyA* gene (Science. 2003; 299(5615): 2074-6.) | atgaaacaatatttagatttactcaatcgcgtattaactgaaggaactgagaaaagtgaccgtacaggaaccggaacgatcagtgtgttcggacatcagatgcgtttcaacctcgacgagggtttcccgtgtctgaccaccaaaaaactgcatctgaaatcaatcatctacgagttactctggtttctgcaaggtgatacgaacgcgaaatatctgcaagaacacggtgtacgcatctggaacgaatgggcggacgagaacggtgatttagggcatatctatggctatcagtggcgttcgtggcccgactacgacggcggattcatcgaccagatcagcgaagcggtagagacgatcaagcacaatcccgactcccgccgtatcattgtcagcgcctggaatgtagccgatttaaagaatatgaacctgcctccctgtcatgccttcttccagttttacgtggcagacggtcggctgagcctgcaactttaccagcgcagcgcggatatatttctcggagtcccgttcaatatcgcatcatatgcactgctgctacaaatgatggcgcaggtgacaggactgaaagcgggtgaatttatccatacacttggcgatgcccatatctatctgaaccacttggatcaggtcaaattgcagcttagccgcgaaccacgcgcattgcctcaaatgaaaattaatccggatgtgaagagtatctatgacttccagttcgaagacttcgaactggtgaactacgacccgcatccacatattgccggaatagtagcggtataa |
| Mutated *thyA* | Mutated *thyA* gene (This work) | atgaaacaatatttagatttactcaatcgcgtattaactgaaggaactgagaaaagtgaccgtacaggaaccggaacgatcagtgtgttcggacatcagatgcgtttcaacctcgacgagggtttcccgtgtctgaccaccaaaaaactgcatctgaaatcaatcatctacgagttactctggtttctgcaaggtgatacgaacgcgaaatatctgcaagaacacggtgtgcgtatttggaatgagtgggcggacgagaacggtgatttagggcatatctatggctatcagtggcgttcgtggcccgactacgacggcggattcatcgaccagatcagcgaagcggtagagacgatcaagcacaatcccgactcccgccgtatcattgtcagcgcctggaatgtagccgatttaaagaatatgaacctgcctccctgtcatgccttcttccagttttacgtggcagacggtcggctgagcctgcaactttaccagcgcagcgcggatatatttctcggagtcccgttcaatatcgcatcatatgcactgctgctacaaatgatggcgcaggtgacaggactgaaagcgggtgaatttatccatacacttggcgatgcccatatctatctgaaccacttggatcaggtcaaattgcagcttagccgcgaaccacgcgcattgcctcaaatgaaaattaatccggatgtgaagagtatctatgacttccagttcgaagacttcgaactggtgaactacgacccgcatccacatattgccggaatagtagcggtataa |
| NBU2 integrase | *intN2* tyrosine integrase gene to integrate plasmid onto the genome of *B. thetaiotaomicron*. (Cell Syst. 2015; 1(1): 62-71) | atgaatatcaagcgcaacatcattttgcattggagagccggaaaaagaacggtgtgccaatcgtagagaacgtacccatccgtatgcgtgtcatctttgccagccaacgcatcgagtttacaacgggctaccggattgacgtagccaaatgggatgcagataagcagcgggtaaagagcggatgtaccaacaagctaaagcaaagtgcagccgaaatcaatacggacttgctgaaatactatgccgaaatccagaatattttcaaggaatttgaggtgcaggaggtcatgccaacgacccaacagttgaaggaagctttcaacatgagaatgaaagacaccagcgaagaacagccggaagaagcccctgtcagcttttgggaggtgttcgatgagtttgtaaaagagtgcggtaaccagaataactggacggcatccacctatgaaaaatttgcagcagtgaggaaccacctcaaagagttcaaggaggatgcaacgttcaactatttcaacgagtttggattgaacgaatacgtcaacttcctgcgtgacaccaaggatatgagaaacagcaccatcggcaagcaaatgggattcctcaaatggttcctgcgctggagcttcaagaaaggacatcatcagaacattgcatacgatacgttcaaaccgaaactgaaaaccacctcgaaaaaagtaatcttcctgacttgggatgaactgaacaagctgaaagactaccagatacccaaggataagcaatacctggaacgtgtgcgtgatgttttcctgttctgc |

| | | |
|---|---|---|
| | | tgctttacgagtttgcggtattcggatgttcgcaatctgaaaagaagcgatgtgaagtccgaccacatcgaaataaccacagtcaagactgccgacagcctgacgattgaactgaacaaatacagcaaagccatactggacaaatacaaggacatccatttcgagaattacatggctctgcccgtcatcagcaaccagaagatgaacgattacctgaaagagctgggcgaactggcagaaatcaacgagcctgtacgggaaacctactacaagggaaatgaacgtattgatgaagtcacacccaaatacgctttgctcagtacccatgcaggaagaaggacattcatctgcaatgcgctggctctcggaatcccggcacaggtggtcatgaaatggacgggacacagcgactacaaagctatgaaaccctacattgacatagcggatgatattaaggcaaatgccatgaacaagtttaatcaacta |
| *attN2* | Recombination site of pNBU2 plasmid with *attBT* sites on the genome of *B. thetaiotaomicron.* (Cell Syst. 2015; 1(1): 62-71) | cctgtctctccgcaaaaaacgct |
| TcR | Tetracycline resistance gene (Genbank: NZ_CYYO01000017) | gtgcgtttcgacaatgcatctactgtagtatattattgcttaatccaaatgaatattataaatttaggaattcttgctcacattgatgcaggaaaaacttccgtaaccgagaatctgctgtttgccagtggagcaacggaaaagtgcggccgtgtggataatggtgacaccataacggactctatggatatagagaaacgtagaggaattactgtccgggcttctacgacatctattatctggaatggagtgaaatgcaatatcattgacactccgggacacatggattttattgcggaagtggagcggacattcaaaatgcttgatggagcagtcctcatcttatccgcaaaggaaggcatacaagcgcagacaaagttgctgttcagtactttacaaaagctgcaaatcccgacaattatatttatcaataagattgaccgtgccggtgtgaatttggagcgtttgtatatggatataaaaacaaatctgtcgcaagatgtcctgtttatgcaaactgttgtcgatggatcggtttatccggtttgctcccaaacatatataaaggaagaatacaaagaatttgtatgcaaccatgacgacgatatattagaacgatatttggcggatagcgaaatttcaccggctgattattggaatacgataatcgctcttgtggcaaaagccaaagtctatccggtgctacatggatcagcaatgttcaatatcggtatcaatgagttgttggacgccatttcttcttttatacttcctccggcatcagtctcaaacagactttcagcttatctctataagatagagcatgaccccaaagggcataaaagaagttttcttaaaataattgacggaagtctgagacttcgagacgttgtaagaatcaacgattcggaaaaattcatcaagattaaaaatctaaagactatttatcagggcagagagataaatgttgatgaagtgggtgccaatgatatcgcgattgtagaagatatagaagattttcgaatcggagattatttaggtgctaaaccttgtttgattcaaggattatctcatcagcatcccgctctcaaatcctccgtccggccaaataagcccgaagagagaagcaaggtgatatccgctctgaatacattgtggattgaagacccgtctttgtccttttccataaactcatatagtgatgaattggaaatctcgttatatggtttgacccaaaaggaaatcatacagacattgctggaagaacgattttccgtaaaggtccattttgatgagatcaagactatctacaaagaacgacctataaaaaaggtcaataagattattcagatcgaagtaccacccaacccttactgggccacaatagggctgactcttgaacccttaccgttaggggcagggttgcaaatcgaaagtgacatctcctatggttatctgaaccattcttttcaaaatgccgtttttgaagggattcgtatgtcttgccaatctggtttacatggatgggaagtgacagatctgaaagtaactttactcaagccgagtattatagcccggtaagtacacctgctgatttcagacagctgaccccttatgtcttcaggctggctttgcaacagtcaggtgtggacattctcgaaccgatgctctgttttgagttgcagataccccaagtagcgagttccaaagctattacagatttgcaaaaactgatgtctgagattgaagacatcagttg |

| | | |
|---|---|---|
| | | taataatgagtggtgtcatattaaagggaaagttccattaaatacaagtaaagactatgcctcagaagtaagttcgtacactaagggcttaggcatttttatggttaagccatgtgggtatcaaataacaaaagacggttattctgataatatccgcatgaacgaaaaagataaacttttattcatgttccaaaaatcaatgtcattaaaataa |
