## Supplementary Table 5 for "Cas9-Assisted Biological Containment of a Genetically Engineered Human Commensal Bacterium and Genetic Elements"

**Supplementary Table 5. List of plasmids used in experiments.**

| Name | Description | Purpose | Figure |
|---|---|---|---|
| pNH9104 | pNBU2-phes-$P_1$-NS-cas9-$P_{BT1311}$-rpiL*-nanoluc | First recombination | 2, S2 |
| pNH9040 | pNBU2-phes-$P_1$-NS-cas9-$P_{BT1311}$-crRCN-rpiL*-nanoluc | First recombination | 2, S2 |
| pNH9058 | pNBU2-phes-$P_1$-NS-cas9-$P_{BT1311}$-crR7N-rpiL*-nanoluc | First recombination | 2, S2 |
| pNH9059 | pNBU2-phes-$P_1$-NS-cas9-$P_{BT1311}$-crR10N-rpiL*-nanoluc | First recombination | 2, S2 |
| pNH9060 | pNBU2-phes-$P_1$-NS-cas9-$P_{BT1311}$-crR12N-rpiL*-nanoluc | First recombination | 2, S2 |
| pNH9106 | pNBU2-phes-$P_1$-taRNA1-cas9-$P_{BT1311}$-crRCN-rpiL*-nanoluc | First recombination | 2, S2, S3 |
| pNH9107 | pNBU2-phes-$P_1$-taRNA2-cas9-$P_{BT1311}$-crRCN-rpiL*-nanoluc | First recombination | 2, S2, S3 |
| pNH9108 | pNBU2-phes-$P_1$-taRNA7-cas9-$P_{BT1311}$-crRCN-rpiL*-nanoluc | First recombination | 2, S2, S3 |
| pNH9109 | pNBU2-phes-$P_1$-taRNA3-cas9-$P_{BT1311}$-crRCN-rpiL*-nanoluc | First recombination | 2, S2, S3 |
| pNH9142 | pNBU2-phes-$P_1$-taRNA4-cas9-$P_{BT1311}$-crRCN-rpiL*-nanoluc | First recombination | 2, S2, S3 |
| pNH9143 | pNBU2-phes-$P_1$-taRNA5-cas9-$P_{BT1311}$-crRCN-rpiL*-nanoluc | First recombination | 2, S2, S3 |
| pNH9144-2 | pNBU2-phes-$P_1$-taRNA6-cas9-$P_{BT1311}$-crRCN-rpiL*-nanoluc | First recombination | 2, S2, S3 |
| pNH9169 | pNBU2-phes-$P_{BT1311}$-NS-cas9-$P_1$-rpiL*-nanoluc | First recombination | 2, 4, S2 |
| pNH9168 | pNBU2-phes-$P_{BT1311}$-NS-cas9-$P_1$-crRCN-rpiL*-nanoluc | First recombination | 2, S2 |
| pNH9188 | pNBU2-phes-$P_{BT1311}$-taRNA6-cas9-$P_1$-crRCN-rpiL*-nanoluc | First recombination, Evaluation of HGT | 2-7, S1-S9 |
| pNH9195 | pNBU2-phes-$P_{cfxA}$-sgRNA1 | Second recombination | 3, S2 |
| pNH9196 | pNBU2-phes-$P_{cepA}$-sgRNA1 | Second recombination | 3, 4, 6, 7, S1, S2, S4, S5, S7, S8, S9 |
| pNH9198 | pNBU2-phes-$P_{cepA}$-NT | Second recombination | 4, 7 S2, S5, S9 |
| pYL100 | pNBU2-NBU2 integrase-*attN*-$P_{cepA}$-sgRNA1-emR | Integration of a plasmid | 4, S6 |

| | | | |
|---|---|---|---|
| pYL114 | pNBU2-NBU2 integrase-*attN*-$P_{cepA}$-NT-emR | Integration of a plasmid | 4, S6 |
| pYL101 | pNBU2-NBU2 integrase-*attN*-$P_{cepA}$-sgRNA12-emR | Integration of a plasmid | 4, S6 |
| pYL103 | pNBU2-NBU2 integrase-*attN*-$P_{cepA}$-sgRNA13-emR | Integration of a plasmid | 4, S6 |
| pYL106 | pNBU2-NBU2 integrase-*attN*-$P_{cepA}$-sgRNA15-emR | Integration of a plasmid | 4, S6 |
| pYL107 | pNBU2-NBU2 integrase-*attN*-$P_{cepA}$-sgRNA16-emR | Integration of a plasmid | 4, S6 |
| pYL112 | pNBU2-NBU2 integrase-*attN*-$P_{cepA}$-sgRNA20-emR | Integration of a plasmid | 4, S6 |
| pNH9223 | pNBU2-phes-*thyA*-emR | Evaluation of HGT | 4, 6, S5, S7 |
| pNH9233 | pNBU2-phes-Mutated *thyA*-emR | Evaluation of HGT | 4, 6, S5 |
| pNH9225 | pNBU2-phes-$P_{BT1311}$-taRNA6-$P_{cfxA}$-sgRNA1-cas9-$P_1$-crRCN-rpiL*-nanoluc | Evaluation of HGT | 5 |
| pNH9226 | pNBU2-phes-$P_{BT1311}$-taRNA6-$P_{cepA}$-sgRNA1-cas9-$P_1$-crRCN-rpiL*-nanoluc | Evaluation of HGT | 5 |
| pNH9227 | pNBU2-phes-$P_{BT1311}$-taRNA6-$P_{cepA}$-NT-cas9-$P_1$-crRCN-rpiL*-nanoluc | Evaluation of HGT | 5 |
| pNH9216 (Cell Syst. 2015; 1(1): 62-71) | pNBU2-NBU2 integrase-*attN*-emR | Integration of a plasmid | 7, S8, S9 |
| pNH9230-2 | pNBU2-NBU2 integrase-*attN*-tcR | Integration of a plasmid | 7, S8 |
