## Supplementary Table 6 for "Cas9-Assisted Biological Containment of a Genetically Engineered Human Commensal Bacterium and Genetic Elements"

**Supplementary Table 6. List of primers used in experiments.**

| Name | Description | Figure | Sequence |
|---|---|---|---|
| Seq-*thyA*-F | PCR identification of horizontal gene transfer of the synthetic gene cassettes. | 5 | cactgtacgcagctaacac |
| mmD663 | PCR identification of horizontal gene transfer of the synthetic gene cassettes. | 5 | atcaccagcgtaccgtaatg |
| qPCR-EmR-F | PCR identification of horizontal gene transfer of the EmR gene. | 5, S7 | aacagcaatgctagtgatctg |
| qPCR-EmR-R | PCR identification of horizontal gene transfer of the EmR gene. | 5, S7 | atttggcagcataccttacaac |
| Rec-5'-*thyA*-F | PCR identification of integration of the synthetic gene cassettes. | S2 | attcattgatttattcagcgccatc |
| Rec-*thyA*-3'-R | PCR identification of integration of the synthetic gene cassettes and the sgRNA1. | S2 | agatgctttagatgagcaaattct g |
| CRI-sgRNA-3'-F | PCR identification of integration of the sgRNA1. | S2 | ggaacgaatggggttttagag |
| CRI-sgRNA5-3'-F | PCR identification of integration of the sgRNA (Non-targeting control). | S2 | gtggattctctagactttacttgag |
| *thyA*-F | PCR identification of horizontal gene transfer of the *thyA* gene. | S5 | tgttcggacatcagatgcgtttc |
| *thyA*-R | PCR identification of horizontal gene transfer of the *thyA* gene. | S5 | ggaagaaggcatgacaggga g |
